# Modular and efficient pre-processing of single-cell RNA-seq

**DOI:** 10.1101/673285

**Authors:** Páll Melsted, A. Sina Booeshaghi, Fan Gao, Eduardo Beltrame, Lambda Lu, Kristján Eldjárn Hjorleifsson, Jase Gehring, Lior Pachter

**Author notes:** Authors contributed equally (Páll Melsted, A. Sina Booeshaghi).

## Abstract

Analysis of single-cell RNA-seq data begins with pre-processing of sequencing reads to generate count matrices. We investigate algorithm choices for the challenges of pre-processing, and describe a workflow that balances efficiency and accuracy. Our workflow is based on the kallisto (https://pachterlab.github.io/kallisto/) and bustools (https://bustools.github.io/) programs, and is near-optimal in speed and memory. The workflow is modular, and we demonstrate its flexibility by showing how it can be used for RNA velocity analyses. Documentation and tutorials for using the kallisto | bus workflow are available at https://www.kallistobus.tools/.

## Introduction

The quantification of transcript or gene abundances in individual cells from a single-cell RNA-seq (scRNA-seq) experiment is a task referred to as pre-processing^1^. The pre-processing steps for scRNA-seq bear some resemblance to those used for bulk RNA-seq^2^ and are in principle straightforward: cDNA reads originating from transcripts must be partitioned into groups according to cells of origin, aligned to reference genomes or transcriptomes to determine molecules of origin, and reads originating from PCR-duplicated molecules must be “collapsed” so they are counted only once during quantification. The collapsing step can be facilitated with unique molecular identifiers (UMIs), which are sequences that serve as barcodes for molecules^3^. The challenges in pre-processing single-cell RNA-seq lie in the tradeoffs that must be considered in determining choices for how, and in which order, to execute the various steps. For example, in droplet-based scRNA-seq protocols, collapsing UMIs to account for PCR duplication can be performed naïvely by associating all reads that align to the same gene, with the same UMI, to a single molecule^4^. This computationally efficient procedure is based on an assumption that all reads with identical UMIs that align to the same gene arise via PCR duplication of a single molecule. Alternatively, this assumption can be relaxed, resulting in a problem formulation that is NP-complete, i.e. computationally intractable to solve optimally^5^. Another challenge in scRNA-seq pre-processing is the amount of data that must be processed. A single cell experiment can generate 10^6^ - 10^10^ reads from 10^3^ - 10^6^ cells^6^. This is leading to bottlenecks in analysis: for example, the current standard program for pre-processing 10x Genomics Chromium scRNA-seq, the Cell Ranger software^7^, requires approximately 22 hours to process 784M reads^5^ using 1.5 Tb of disk space (Supplementary Table 1). For this reason, a number of new, faster workflows for scRNA-seq pre-processing based on pseudoalignment^8^ have recently been developed^5,9^. However, despite improvements in running time, current workflows have memory requirements that increase with data size^5^, a situation that is untenable given the pace of improvement in technology and the corresponding increase in data volume.

In recent work we introduced a new format for single-cell RNA-seq data that makes possible the development of efficient workflows by virtue of decoupling the computationally demanding step of associating reads to transcripts and genes (alignment), from the other steps required for scRNA-seq pre-processing^10^. The new format, called BUS (Barcode, UMI, Set), can be produced by pseudoalignment, and rapidly manipulated by a suite of tools called BUStools. To illustrate the utility, efficiency, and flexibility of this approach for scRNA-seq pre-processing, we describe a Chromium pre-processing workflow based on reasoned choices for the key pre-processing steps. While we focus on Chromium, our workflow is general and can be used with other technologies. We show that our pre-processing workflow is much faster and more memory efficient than existing methods, and we demonstrate the power of modular processing with the BUS format by developing a fast RNA velocity analysis workflow^11^. We also validate the design decisions underlying the Cell Ranger workflow. Our benchmarking and testing is comprehensive, comprising analysis of almost two dozen datasets and greatly eclipsing the scale of testing that has been performed for current workflows.

## Results

In designing a scRNA-seq pre-processing workflow, we began by investigating each required step: correction of barcodes, collapsing of UMIs, and assignment of reads to genes. To achieve single-cell resolution, the Chromium technology produces barcode sequences that are used to associate cDNA reads with cells, and we began by considering the efficiency-accuracy tradeoffs involved in grouping reads with the same, or similar, barcodes to define the contents of individual cells. The Chromium barcodes arise from a “whitelist”, a set of pre-defined sequences that are included with the Cell Ranger software. Grouping reads according to barcode is therefore straightforward, except for the fact that barcodes may contain sequencing errors. The Cell Ranger workflow corrects all barcodes that are one base-pair change away (Hamming distance 1) from barcodes in the whitelist. An examination of a benchmark panel of 20 datasets revealed that this error correction approach can be expected to rescue, on average, 0.8% of the reads in an experiment (Figure 1a,b), a calculation based on an inferred error rate per base for each dataset (Methods, Supplementary Table 1). Thus, correction of barcodes Hamming distance 2 away from whitelist barcodes would rescue, on average, a negligible number (0.0038%) of reads (Methods). We therefore implemented a Hamming distance 1 correction method in our workflow via the **bustools correct** command.

**Figure 1:**
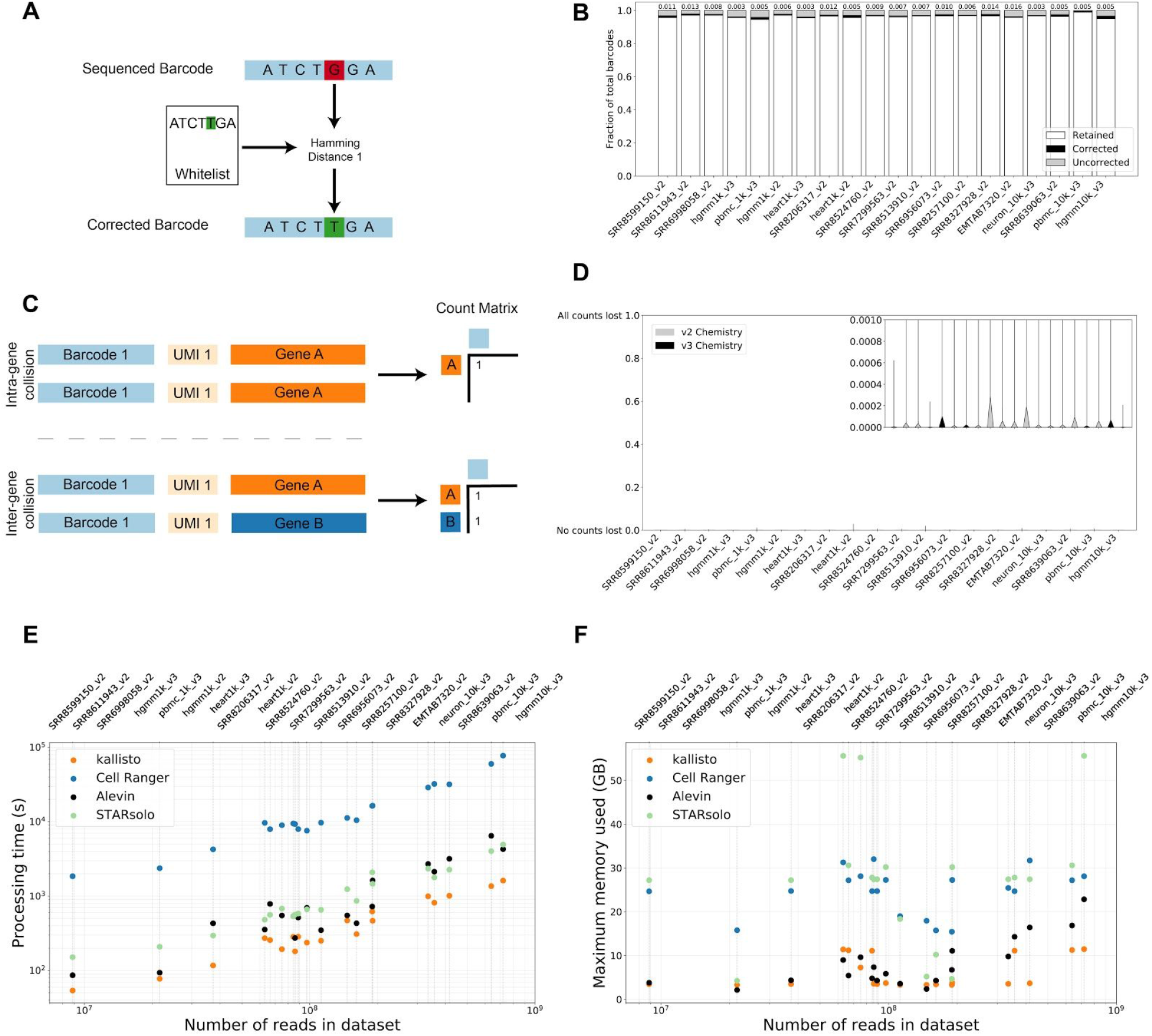
(A) Error correction of barcodes 1 mismatch away from barcodes in a whitelist. (B) Analysis of barcode fidelity in the benchmark panel (20 datasets) showing barcodes matching the whitelist (“retained”, white), barcodes that are Hamming distance 1 away from the whitelist that were corrected (black) and uncorrected barcodes (gray). (C) UMI collapsing within genes. Fraction of UMIs lost per gene across cells in the benchmark panel due to over-collapsing. Running time of kallisto (orange), Cell Ranger (blue), Alevin (black), and STARsolo (green) for pre-processing the benchmark panel. (F) Memory usage of kallisto (orange), Cell Ranger (blue), Alevin (black) and STARsolo (green) for pre-processing the benchmark panel.

Next, in considering how to collapse UMIs, we first investigated the extent to which “collisions” occur, i.e. cases where the same UMI occurs in reads originating from two different molecules^12^. While inter-gene collisions can be directly measured, intra-gene collisions cannot be distinguished from PCR duplicates. To estimate the intra-gene collision rate we first calculated, for each cell in the benchmark panel, the effective number of UMIs in each of the associated droplets (Supplementary Figure 1, Supplementary Note). This estimate, along with the number of inter-gene collisions and distinct UMIs observed, allowed us to estimate the extent of intra-gene collision, and therefore the count loss due to naïve collapsing of UMIs by gene (Methods, Supplementary Note). We found for the benchmark panel that the average percentage of lost counts per gene per cell due to naïve collapsing was less than 0.003% for v2 chemistry and 0.000048% for v3 chemistry (Figure 1c,d). Thus, we decided to apply naïve collapsing as it is computationally efficient and effective based on empirical evidence. This was implemented in the **bustools count** command. Notably, the recently published Alevin collapsing algorithm^5^ will overestimate gene counts because reads with the same UMI pseudoaligned to the same gene are very likely to be from the same molecule even if they pseudoalign to distinct transcripts. In fact, our analysis suggests such situations result from missing or incorrect annotation^13^ rather than from collisions of two distinct molecules labeled with the same UMI.

One implication of the UMI collapsing analysis is that UMI error correction is possible because UMIs with only one base-pair change away from an abundant UMI are likely to have resulted from sequencing error. To examine the benefit of such a correction we computed the expected number of UMIs that would be corrected with Hamming distance 1 correction, and found that for 10bp and 12bp UMIs only 0.5% and 0.6% of reads would be recovered, respectively, at the error rates observed in published datasets (Methods, Supplementary Figure 2). Moreover, such error correction would require identification of abundant UMIs in lieu of a whitelist, adding time and complexity to the workflow. While we believe such error correction may be warranted in the case of longer UMIs (Supplementary Figure 2), we did not include it in our workflow.

In most scRNA-seq pre-processing workflows, assignment of cDNAs to genes utilizes genome alignment^4,14,15^. Since detailed base-pair alignment is not necessary to generate a count matrix, pseudoalignment to a reference transcriptome^8^ suffices. Moreover, pseudoalignment has been shown to be highly concordant with alignment for the purposes of quantification in bulk RNA-seq^16^. To test this hypothesis we compared counts obtained by pseudoalignment using the kallisto program^8^ with counts produced via Cell Ranger which is based on the STAR aligner^17^. Analysis of an *Arabidopsis thaliania* scRNA-seq dataset (Figure 2), confirms that there is a high correlation between pseudoalignment and alignment based counting, however in one dataset (pbmc10k_v3, Supplementary Figure 3.19) we found that pseudoalignment produced more counts than alignment. Specifically, in the FGF23 (ENSG00000118972) gene, Cell Ranger had many fewer counts than kallisto. We hypothesized that the reason for this discrepancy was the presence of reads from unspliced transcripts crossing splice junction boundaries, and therefore being erroneously pseudoaligned to the transcriptome. To test this we created a modified index that included a 90 base-pair overlap into the exon and the intron (one base-pair less than the length of the reads) to capture such reads and confirmed that it resolved the discrepancy (Supplementary Figure 4). We observed this problem to be rare and therefore did not deem it to be essential in a standard processing workflow. It may be that consideration of such reads will be crucial for nuclear scRNA-seq analyses^18^, when the abundance of such intronic junction reads will be problematic for naïve pseudoalignment.

**Figure 2:**
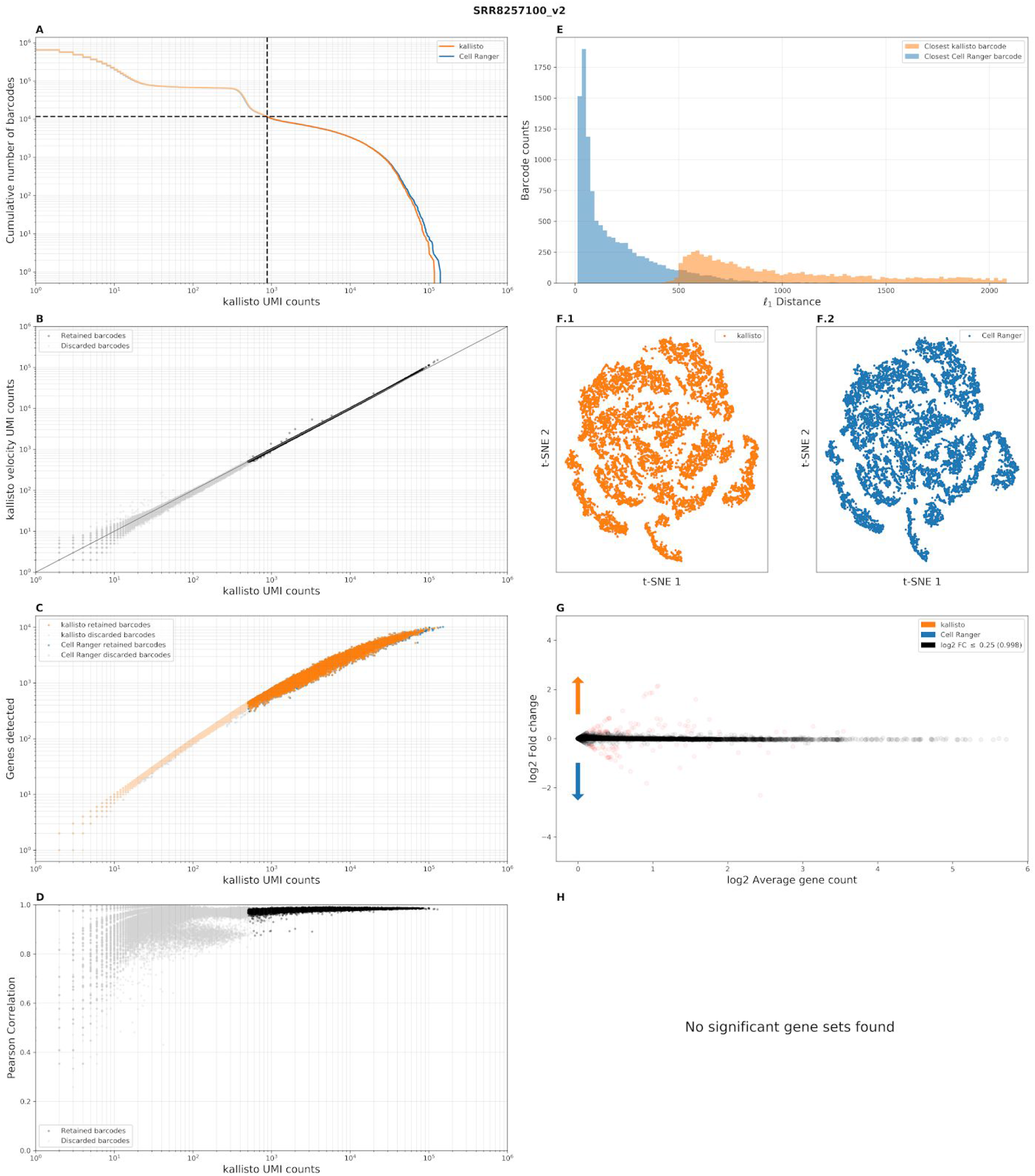
Benchmark panel of single-cell RNA-seq data from *Arabidopsis thaliana*, Ryu et al. 2019^19^ (SRR8257100). Darker points\lines are retained barcodes and lighter points\lines are discarded barcodes. (A) “Knee plots” for kallisto and Cell Ranger showing, for a given UMI count (x-axis), the number of cells that contain at least that many UMI counts (y-axis). The dashed lines correspond to the Cell Ranger filtered cells. (B) Correspondence in the number of distinct UMIs per cell between the workflows. (C) Genes detected by kallisto and Cell Ranger as a function of distinct UMI counts per cell. (D) Pearson correlation between gene counts as a function of the distinct UMI counts per cell. (E) The *l*_1_ distance between gene abundances for each kallisto cell and its corresponding Cell Ranger cell (blue) and the *l*_1_ distance between the gene abundances for each kallisto cell and the closest kallisto cell (orange). (F) kallisto t-SNE from the first 10 principal components. (G) Cell Ranger t-SNE from the first 10 principal components. (H) Significant differential gene sets between Cell Ranger and kallisto.

Thus, our workflow consists of pseudoalignment of reads to a reference transcriptome to generate a BUS (barcode, UMI, set) file, and subsequent processing to correct barcode errors and produce a count matrix (Supplementary Figure 5). To ensure that memory usage is constant in the number of reads, the BUS files are sorted by barcode prior to counting using the **bustools sort** command. While this workflow is very similar to that of Cell Ranger, it is not identical. Since Cell Ranger is widely used, we investigated the extent to which the Cell Ranger results are concordant with our workflow. We processed 20 datasets (Supplementary Table 1), chosen to contain a range of reads depths (from 8,860,361 to 721,180,737 reads per sample and 2,243 to 201,952 reads per cell) and to represent scRNA-seq from a range of tissues and species (*Arabidopsis thaliania*^19^, *Caenorhabditis elegans*^20^, *Danio rerio*^21^, *Drosophila melanogaster*^22^, *Homo sapiens*^23,24^, *Mus musculus*^25–29^, *Rattus norvegicus*^29^). We found a high degree of concordance with respect to quality control metrics (Figure 2a-h, Supplementary Figure 3). Crucially, in all datasets, in a joint analysis of kallisto and Cell Ranger counts, the closest cell to a kallisto cell was its associated Cell Ranger cell, i.e. the Cell Ranger cell with the same barcode sequence. Furthermore, gene count correlations between individual cells passing Cell Ranger filtering criteria were almost always above 90%, and frequently as high as 99%.

To assess the extent to which differences between Cell Ranger and kallisto affect biological inferences, we also compared Cell Ranger to kallisto in a variety of typical downstream analyses on the 10x Genomics E18 mouse 10k brain cells dataset. The Cell Ranger analysis produces structures similar to that of kallisto when projected to the first two principal components (PCs) and to two dimensions of tSNE (Supplementary Figure 6.1). The results of Leiden clustering^30^ are similar regardless of pre-processing workflow, though while most clusters can be uniquely matched between Cell Ranger and kallisto, there are a few cases of cluster merging and splitting (Supplementary Figure 6.2). We performed differential expression (DE) analysis to identify marker genes of the clusters, and then performed gene set enrichment analysis (GSEA) on the marker genes for cell type annotation (Supplementary Figure 6.3). The marker genes and their corresponding gene sets were highly correlated between the workflows. In both Cell Ranger and kallisto results, most clusters are neuronal, and the clusters for erythrocytes (cluster 16 in both), endothelial cells (cluster 21 in kallisto, cluster 19 in Cell Ranger), and immune cells (clusters 20 and 22 in kallisto cluster 17 in Cell Ranger) can be clearly identified based on marker genes (Supplementary Figure 6.3). Correlation between the same barcodes in kallisto and Cell Ranger with the top cluster marker genes is very high, with both the Pearson and Spearman correlation coefficient above 0.9 for the vast majority of cells (Supplementary Figure 6.4). Pseudotime inference with Cell Ranger and kallisto resulted in concordant trajectories from neuronal precursor cells to two populations of neurons, with the same trajectory topology and similar pseudotime values along the trajectory (Supplementary Figure 6.5). In a separate mixed species dataset, the number and proportion of UMIs from human and mouse cells are similar between Cell Ranger and kallisto (Supplementary Figure 7). Overall, these results suggest that the Cell Ranger workflow produces results consistent with our method, not only at the level of dataset summary statistics, but also in downstream analyses.

The modularity of our approach makes possible the rapid implementation of alternative workflows. To illustrate this we developed an RNA velocity workflow. By including intron sequences in the index for pseudoalignment we were able to identify reads originating from unspliced transcripts, and, using the **bustools capture** command, efficiently created the spliced and unspliced matrices needed for RNA velocity. Our RNA velocity workflow, which is 13 times faster than velocyto^11^ analysis of the same dataset, is suitable for large datasets that were previously challenging to pre-process. To illustrate this we computed RNA velocity vectors for recently published data from the developing mouse retina^31^ consisting of 113,917 cells (Figure 3). We found that six pseudotime marker genes highlighted in Clark et al. 2019^32^ (Crx, Nrl, Otx2, Pax6, Rbpms, Rlbp1) displayed patterns consistent with the RNA velocity vectors, and with the pseudotime analysis of Clark et al.^32^ (Supplementary Figure 8). The velocity analysis reveals new information, namely it identifies developmental states when RNA velocity is changing (Supplementary Figure 8 middle column). We verified the fidelity of our workflow by computing RNA velocity vectors on a dataset from La Manno et al. 2018^11^ and comparing our results to those of the paper (Methods, Supplementary Figure 9). Furthermore, the spliced count matrix agreed with the count matrix obtained in our standard gene expression workflow (Supplementary Figure 10). Despite identifying many more unspliced counts (Supplementary Figure 11), our resultant velocity figure was concordant with that of La Manno et al.^11^.

**Figure 3:**
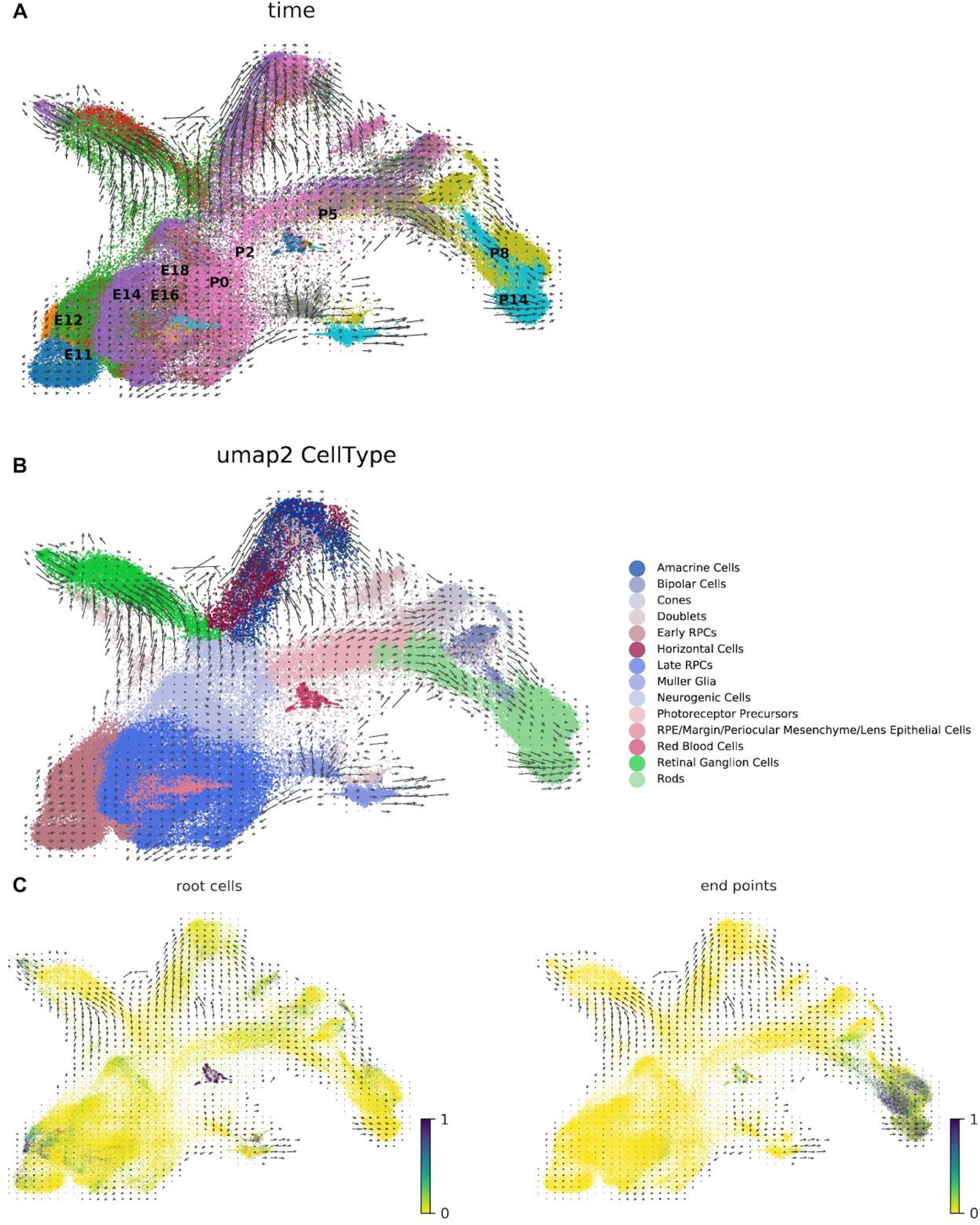
(A) A kallisto and bustools based RNA velocity analysis of the ten stage scRNA-seq retina neurogenesis data from Clark et al. 2019^32^. (B) Clusters annotated by cell type according to Clark et al. (C) Markov diffusion process analysis highlighting source and sink cells and demonstrating that the velocity vector field is consistent with the cells’ developmental trajectory.

## Discussion

Our scRNA-seq workflow is up to 51 times faster than Cell Ranger and up to 4.75 times faster than Alevin. It is also up to 3.5 times faster than STARsolo: a recent version of the STAR aligner adapted for scRNA-seq (Figure 1e, Supplementary Table 2). Importantly, unlike these other programs, our workflow requires a small fixed amount of constant memory that is independent of the number of reads being pre-processed (Figure 1f). It is therefore suitable for low-cost and environmentally conscious cloud computing. Moreover, the workflow is scalable and can in principle be used for pre-processing arbitrary numbers of reads. In benchmarks on the panel described in this paper, kallisto’s running time was comparable to that of the word count (**wc**) command applied to the FASTQ files, suggesting that kallisto is near-optimal in efficiency (Supplementary Figure 12). Our speed and constant memory requirements make RNA velocity tractable for datasets of any size for the first time.

Our UMI collapsing analysis suggests that UMI sequences can be short; even just 5 base-pairs of sequence suffice for identifying molecules thanks to the cell barcode and gene identification for each read serving as auxiliary barcodes (Supplementary Figure 13). Furthermore, the fact that identical UMIs associated with distinct reads from the same gene are almost certainly reads from the same molecule (Figure 1d), makes it possible, in principle, to design efficient assignment algorithms for multi-mapping reads. Reads could be assigned with an expectation-maximization algorithm which is based on estimating the copy number of each molecule in the library using a model as described in the Supplementary Note, and this is a promising direction for future work. An initial attempt at such assignment^5^ appears to improve concordance between single-cell RNA-seq gene abundance estimates and those from bulk RNA-seq. Importantly, the current implementation of our approach can produce transcript compatibility counts which have information about read ambiguity prior to assignment of multi-mapping reads, and can therefore be used to identify isoform-specific changes across cells and cell clusters ^33^.

While we have focused on a workflow for 10x Chromium data, the bustoools commands we implemented are generic and will work with any BUS file, generated with data from any scRNA-seq technology. Distinct technology encodes barcode and UMI information differently, but the **kallisto bus** command can accept custom formatting rules. While the pre-processing steps for error correction and counting may need to be optimized for the distinguishing characteristics of different technologies, the modularity of the bustools based workflow makes such customization possible and easy.

## Supporting information

Supplementary Table 1 Benchmark Panel Summary

Supplementary Table 2 Time & Memory

## Acknowledgments

We thank Vasilis Ntranos and Valentine Svensson for helpful suggestions and comments. We thank Jeff Farrell for the *Danio rerio* gene annotation used to process SRR6956073, John Schiefelbein for the *Arabidopsis thaliana* gene annotation used to process SRR8257100, Justin Fear the *Drosophila melanogaster* gene annotation used to process SRR8513910, and Junhyong Kim and Qin Zhu for the *Caenorhabditis elegans* gene annotation used to process SRR8611943. We thank Julien Roux for suggesting the analysis in Supplementary Figure 10. The benchmarking work was made possible, in part, thanks to support from the Caltech Bioinformatics Resource Center.

## Author Contributions

PM developed the algorithms for bustools and wrote the software. ASB conceived of and performed the UMI and barcode calculations motivating the algorithms. FG implemented and performed the benchmarking procedure, and curated indices for the datasets. ASB and EB designed and produced the comparisons between Cell Ranger and kallisto. LL investigated in detail the performance of different workflows on the 10k mouse neuron data and produced the analysis of that dataset. ASB designed the RNA velocity workflow and performed the RNA velocity analyses. KH developed and investigated the effect of, and optimal choice for, reference transcriptome sequences for pseudoalignment. JG interpreted results and helped to supervise the research. ASB planned, organized and made figures. ASB, EB, PM and LP planned the manuscript. ASB and LP wrote the manuscript.

## Methods

### Benchmark panel data

A diverse set of 20 datasets was compiled for the purpose of benchmarking pre-processing workflows. Datasets produced and distributed by 10x Genomics were downloaded from the 10x Genomics data downloads page: https://support.10xgenomics.com/single-cell-gene-expression/datasets. Six v3 chemistry datasets and two v2 chemistry datasets were downloaded and processed (Supplementary Table 1). Another 12 datasets were obtained from either the SRA or the ENA; all were produced with 10x Genomics v2 chemistry. For six of the datasets (SRR6956073, SRR6998058, SRR7299563, SRR8206317, SRR8327928, SRR8524760) the BAM files were downloaded and the Cell Ranger utility bamtofastq was run to produce fastq files for pre-processing from Cell Ranger structured BAM files. FASTQ files were downloaded directly for the datasets EMTAB7320, SRR8257100, SRR8513910, SRR8599150, SRR8611943, SRR8639063.

Details of all datasets and their accession numbers can be found in Supplementary Table 1.

### Software

The software versions used for the results in the paper were: bustools v0.39.1, Cell Ranger v3.0.0, kallisto v0.46.0, python 3.7, R v3.5.2, Salmon v0.13.1, Scanpy v1.4.1, scvelo 0.1.17, Seurat v3.0, snakemake v5.3.0, STARsolo v2.7.0e, velocyto v0.17.17, wc v8.22 (GNU coreutils), and zcat v1.5 (gzip). All programs were run with default options unless otherwise specified. The code to reproduce this paper is available at https://github.com/pachterlab/MBGBLHGP_2019, kallisto is available at https://pachterlab.github.io/kallisto/ and bustools is available at https://bustools.github.io/. Documentation and tutorials for using the kallisto | bus single-cell RNA-seq workflow are available at https://www.kallistobus.tools/.

### Hardware

All the benchmarks were carried out on a Supermicro server computer (2xXeon® Gold 6152 22-Core 2.1, 3.7GHz Turbo, 12 × 64GB Quad-Rank DDR4 2666MHz memory, 16 × 12TB Ultrastar He12 HUH721212ALE600, 7200 RPM, SATA 6Gb/s HDD) with CentOS7 operating system installed. The running time of all programs were evaluated using eight threads.

### Transcriptome indices

Reference transcriptomes were constructed by processing datasets with Cell Ranger, downloading the constructed Cell Ranger GTF file, and then producing a transcriptome from it and the relevant genome using GFFread (http://cole-trapnell-lab.github.io/cufflinks/file_formats/#the-gffread-utility).

### Inference of per-base sequencing error rate and correctable barcodes

For each dataset, the per-base error rate *p* was estimated by the formula

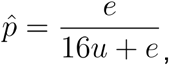

where *u* was the number of barcodes matching the whitelist, and *e* the number of barcodes hamming distance 1 away from a whitelist barcode. This was derived by solving for *p* from the equations

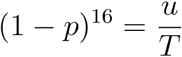

and

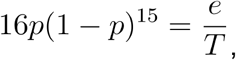

where *T* is the (effective) total number of barcodes.

To estimate the proportion of barcodes Hamming distance 1 away from a whitelist barcode, we computed

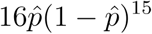

for each dataset, using the estimated per-base error rate 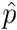.

### UMI collision estimates

A UMI associated with a read from a cell is said to have “collided” if it appears in two or more reads originating from different molecules. To estimate UMI collision rates, two types of information were used. First, reads in the same cell that originate from different genes must have originated from different molecules, and therefore the sharing of a UMI between two such reads was used as an indicator of a collision. Second, for each gene, the number of distinct UMIs associated within it was measured from the data. Based on the assumption that UMIs were sampled uniformly at random from beads, this data was used to estimate the number of intra-gene collisions (see Supplementary Note). The assumption was verified by examining the distribution of UMI counts across cells; the empirical distribution was near-uniform with the exception of a handful of UMIs (Supplementary Note Figure 2).

### Comparative analysis of the benchmark panel datasets

The benchmark panel datasets (Supplementary Table 1) were processed uniformly as follows:

For each dataset a “knee plot”^34^ was constructed for both Cell Ranger and kallisto by plotting, for each cell, the number of distinct UMIs in the cell vs. the number of barcodes with at least that number of UMIs. Then, the distinct number of UMIs for kallisto and Cell Ranger were plotted against each other. Subsequently, for each cell, the number of distinct UMIs was plotted against the number of genes detected. Finally, the Pearson correlation was computed between the gene counts of kallisto and Cell Ranger for each cell.

To investigate the similarity of Cell Ranger to kallisto, the *l*_1_ distance between each corresponding kallisto and Cell Ranger cell was computed. The distance to the nearest kallisto cell was also measured. To visualize the Cell Ranger and kallisto count matrices, t-SNE was performed on the data projected to the 10 principal components computed for each dataset using the opentSNE package (https://github.com/pavlin-policar/openTSNE) with perplexity=30, metric=”Euclidean”, random_state=42, n_iter=750.

To check for systematic differences in the quantification of certain genes between Cell Ranger and kallisto, a differential expression analysis was performed on the matrices produced by the two workflows. First, the matrices were concatenated using the genes determined to be expressed in both methods. Then the counts were normalized using Seurat. DE was performed with logistic regression. Next GSEA with the R package EGSEA on all marker genes with adjusted p-value less than 0.05, to identify classes of genes more likely to be affected by the different workflows. KEGG pathways were used as gene sets.

### Comparative analysis of the 10x Genomics E18 Mouse dataset

Analysis of the Cell Ranger and kallisto pre-processed datasets was performed in R. The DropletUtils package was used to remove empty droplets from the kallisto gene count matrix. For Cell Ranger, the filtered matrix was used. After filtering, genes not detected in any remaining Cell Ranger or kallisto barcode were removed. Seurat was used for basic analysis. First, data was normalized by dividing the UMI count of each gene in each cell by the total UMI counts of that cell, multiplied this number by 10000. Then a pseudocount of 1 was added, and the natural log transform was applied. Subsequently, the normalized data was scaled so the distribution of the expression of each gene would have mean of 0 and standard deviation of 1. Subsequently, 3,000 highly variable genes were selected with the vst method in Seurat. Then principal component analysis was performed on the highly variable genes in the scaled data with the R package irlba called by Seurat. The first 40 principal components were used for tSNE, which was done with the R package Rtsne called by Seurat. Clustering was performed with the Leiden algorithm ^30^ on the kallisto and Cell Ranger matrices. The clustering parameters were 20 nearest neighbors and resolution 1. Differential expression analysis was performed with the logistic regression method described in Ntranos et al.^33^ as implemented in Seurat and applied to the normalized (unscaled) data. Spearman and Pearson correlations were computed for the top 15 cluster marker genes. Gene Set Enrichment Analysis(GSEA) was performed on the top 20 cluster marker genes using the R package EGSEA^35^ with the KEGG pathway gene sets. SingleR^36^ was used to annotate cell types based on correlation profiles with bulk RNA-seq from^37^. Then, the neuronal cell types were used for pseudotime analysis. Pseudotime analysis was done with slingshot^38^ via the Docker container from dyno^39^.

### Species mixing

The 10x Genomics 10k 1:1 Mixture of Fresh Frozen Human (HEK293T) and Mouse (NIH3T3) Cells dataset was analyzed with kallisto and Cell Ranger for the purpose of comparing the resultant banyard plots^40^. Human and mouse genes were identified with their ENSEMBL identifiers. The total number of UMIs mapped to the human and mouse genes in each barcode was calculated with the unfiltered matrices. In Supplementary Figure 7b,c only barcodes present in both the kallisto and Cell Ranger unfiltered matrices were used.

### RNA velocity

A human reference transcriptome FASTA file of exonic transcripts and a reference genome fasta were obtained from the UCSC Genome Browser, build Dec. 2013 GRCh38. A BED file of intronic transcripts, with an (= read length) flanking sequence added to each end, was also obtained from the UCSC Genome Browser. A unique number was appended to the end of each intronic transcript in the BED file. The genome fasta and the intronic BED file were used with **bedtools getfasta** to construct an intronic fasta file. The intronic and exonic fasta files were combined and an index was built with **kallisto index**. The reads were aligned to the index using **kallisto bus**. The barcodes in the resultant BUS file were error corrected with **bustools correct** and then sorted with **bustools sort**. To isolate the intronic counts and exonic counts for each barcode, **bustools capture** was ran twice: once using the list of intronic transcripts and once using the list of exonic transcripts. The spliced count matrices were made by using **bustools count** on the intron-captured split.bus file, and the unspliced count matrices were made by using **bustools count** on the exon-captured split.bus file. Both matrices were loaded into an annotated data frame in a jupyter notebook for downstream analysis.

To perform the comparison to the La Manno et al. 2018 dataset (Supplementary Figures 9,10), the data was first downloaded from the SRA (SRP129388). The cell barcodes were filtered by those in La Manno et al. 2018^11^. The data was pre-processed with the kallisto | bustools workflow. The matrices were loaded into python. The cluster labels were then transferred from La Manno et al. 2018. The velocyto notebook provided with the paper was used to reproduce the results based on the Cell Ranger\Velocyto matrices and the kallisto | bustools matrices.

For the Clark et al. 2019 RNA velocity analysis (Figure 3, Supplementary Figure 8), the data was downloaded from the SRA (GSE118614). First the cell barcodes were filtered by those in Clark et al. 2019. The cluster labels were then transferred from Clark et al. 2019 ^32^ and the standard velocity pipeline from scvelo was run using the kallisto spliced and unspliced matrices.

