## Supplementary Figures for "Modular and efficient pre-processing of single-cell RNA-seq"

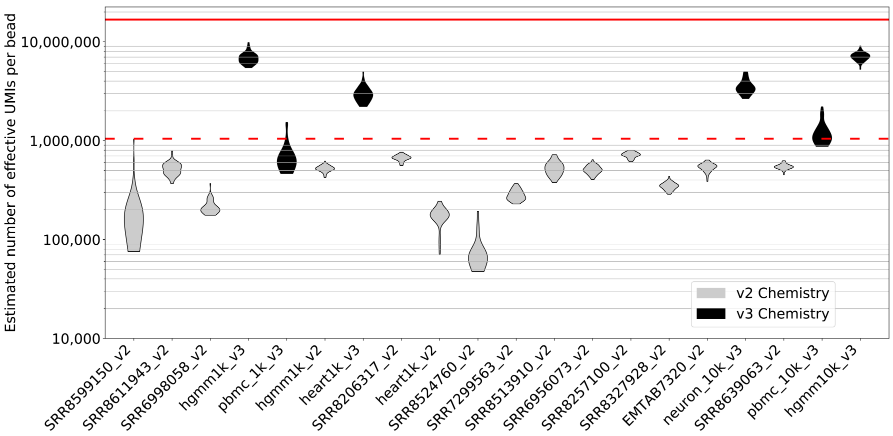


Estimates of the effective number of UMIs per bead for each of the benchmark panel datasets, determined from observed collisions of UMIs across unique genes and assuming UMIs are sampled uniformly with replacement (see Supplementary Note for further details). The dashed red line is the theoretical maximum for the number of UMIs on a v2 chemistry bead ($4^{10} = 1,048,576$) and the red line is the theoretical maximum for the number of UMIs on a v3 chemistry bead ($4^{12} = 16,777,216$). The datasets are ordered by number of reads. UMI pools from 10x Chromium v2 and v3 chemistry are found to be highly complex, with the effective number of UMIs approaching the theoretical maximum in many cases. Our estimates for UMI complexity vary across experiments; this could be due to batch effects, or model misspecification. Sequencing chimeras could also affect UMI complexity estimates, specifically estimates would be increased with more chimeras. This would reduce the estimates of intra-gene collisions due to naïve collapsing.

#### Supplementary Figure 2

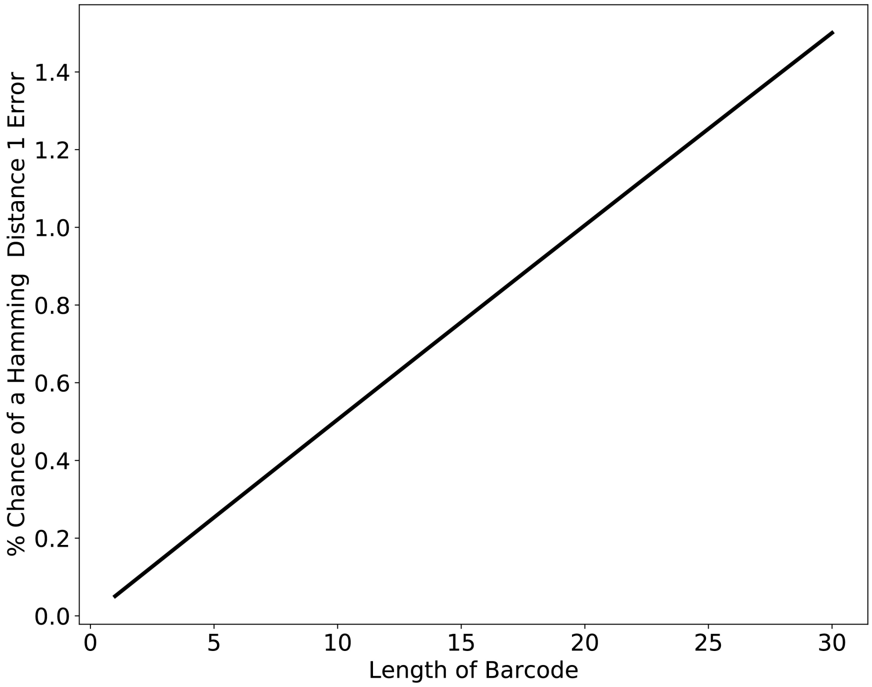


The expected percentage of Barcodes (or UMIs) that will have one error and can therefore be corrected with a Hamming distance 1 correction algorithm. The y-axis displays the value of the function $f(L)$ where $f(L) = 100 \cdot L\hat{p}(1-\hat{p})^{L-1}$, and where $\hat{p}$ is the per base sequencing error probability estimated by averaging the observed error estimates across all the datasets in the benchmark panel (Supplementary Table 1).

#### Supplementary Figure 3

All Supplementary Figure 3 benchmark panels are configured as follows:

(A) "Knee plots" for kallisto and Cell Ranger showing, for a given UMI count (x-axis), the number of cells that contain at least that many UMI counts (y-axis). The dashed lines correspond to the Cell Ranger filtered cells.

(B) Correspondence in the number of distinct UMIs per cell between the kallisto and Cell Ranger.

(C) Genes detected by kallisto and Cell Ranger as a function of distinct UMI counts per cell.

(D) Pearson correlation between gene counts as a function of the distinct UMI counts per cell.

(E) The $l_1$ distance between gene abundances for each kallisto cell and its corresponding Cell Ranger cell (blue) and the $l_1$ distance between the gene abundances for each kallisto cell and the closest kallisto cell (orange).

(F) **1.** kallisto t-SNE from the first 10 principal components. **2.** Cell Ranger t-SNE from the first 10 principal components.

(G) MA plot. Each dot is a gene and the y-axis represents the log2 fold change between the average count for that gene across all cells between kallisto and kallisto velocity (spliced) and the x-axis is the average count for that gene between kallisto and kallisto velocity (spliced).

(H) Significant differential gene sets between Cell Ranger and kallisto.

The darker points correspond to retained barcodes and the lighter points correspond to discarded barcodes.

### Supplementary Figure 3.1

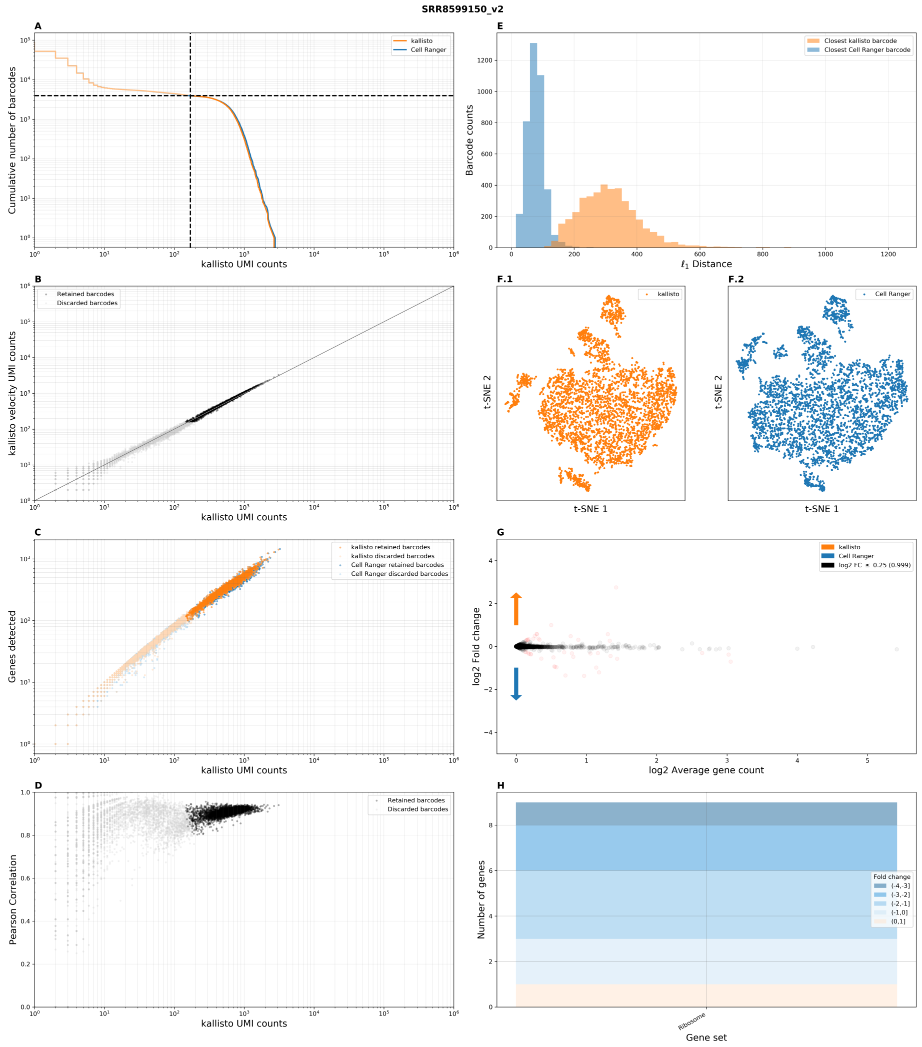


Benchmark panel of dataset `SRR8599150` from O'Koren *et al.* 2019.

### Supplementary Figure 3.2

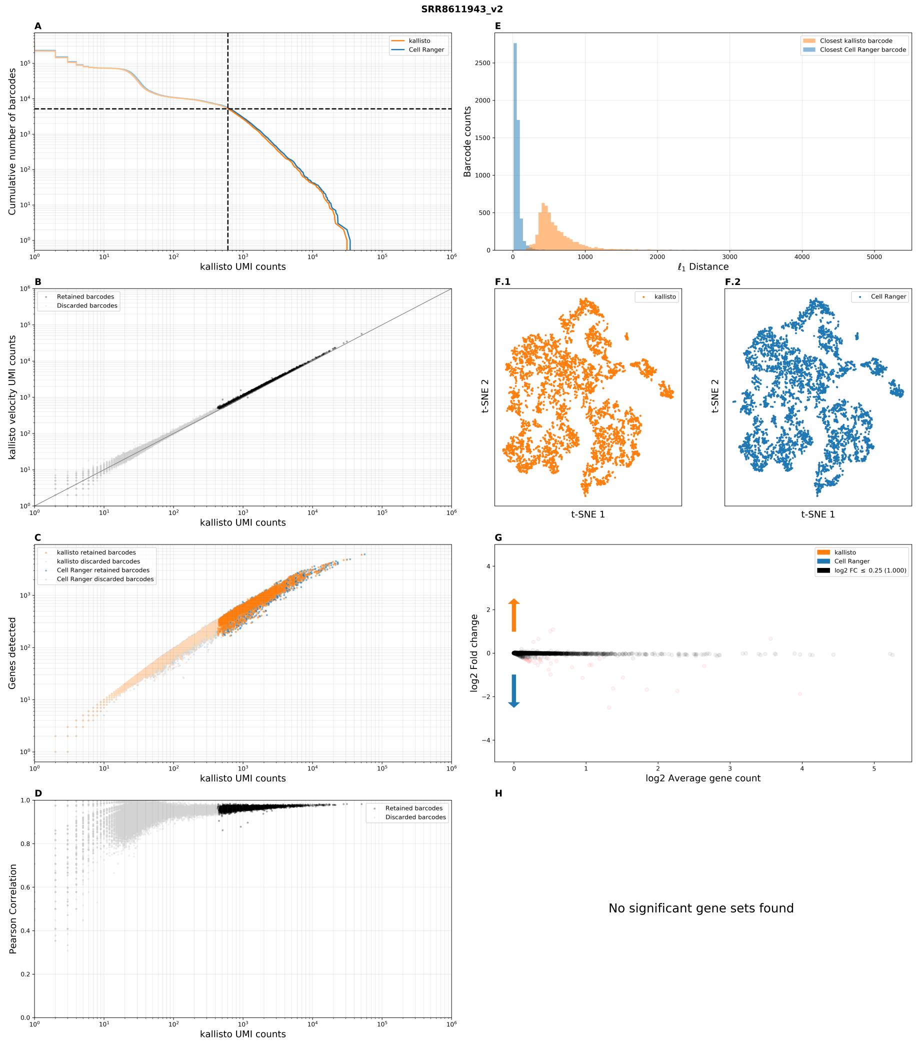


Benchmark panel of dataset SRR8611943 from Packer *et al.* 2019.

### Supplementary Figure 3.3

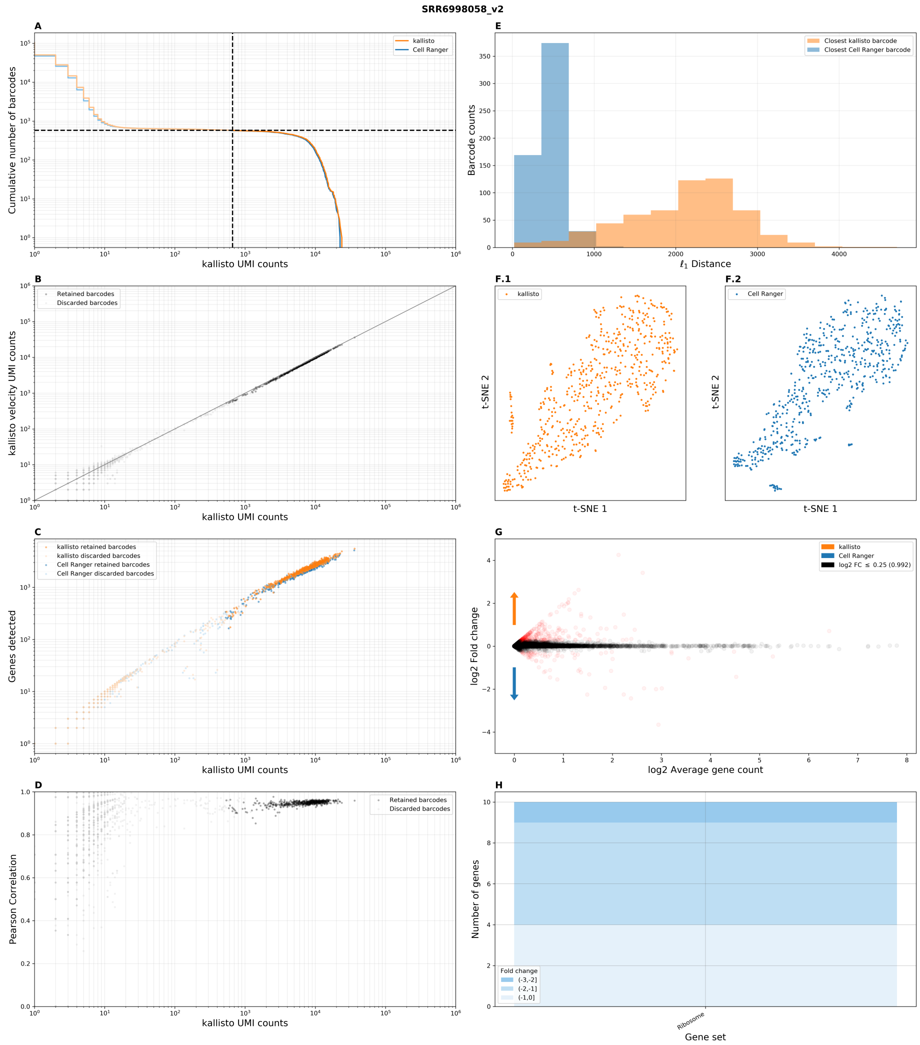


Benchmark panel of dataset `SRR6998058` from Jin *et al.* 2018.

### Supplementary Figure 3.4

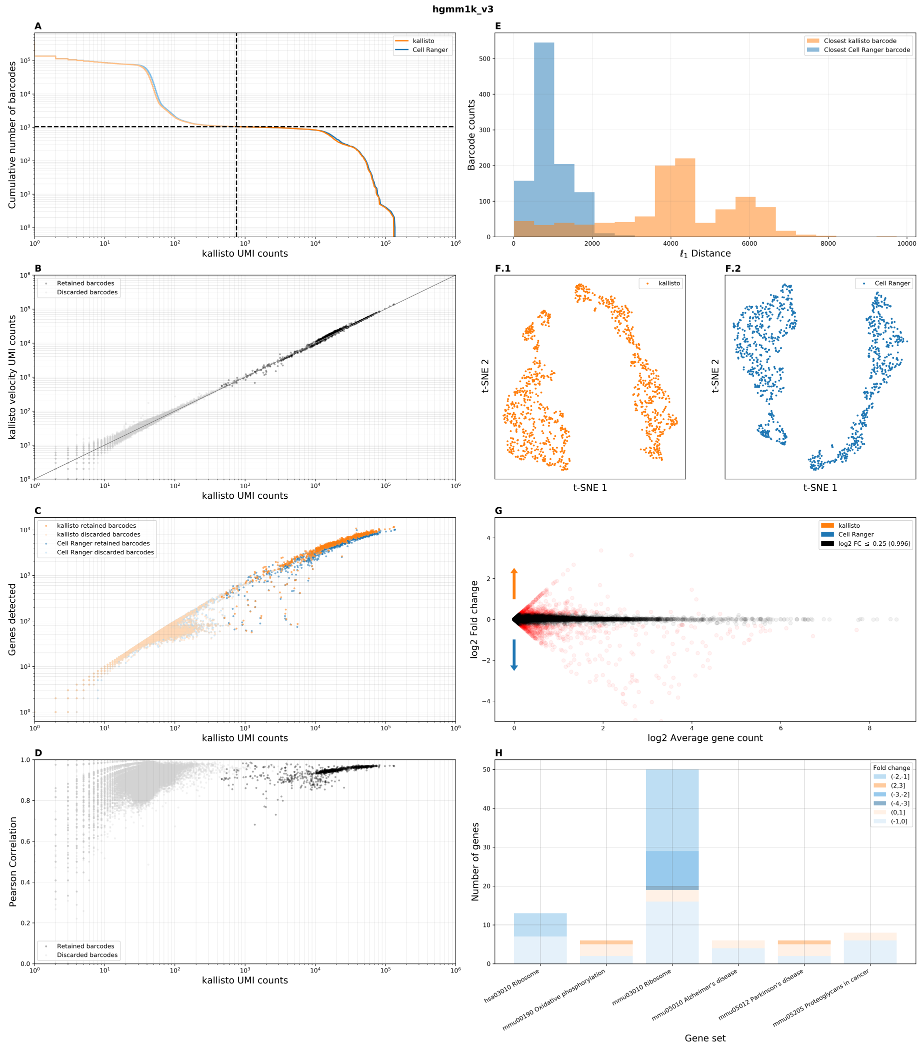


Benchmark panel of dataset `hgmm1k_v3` from 10x Genomics.

### Supplementary Figure 3.5

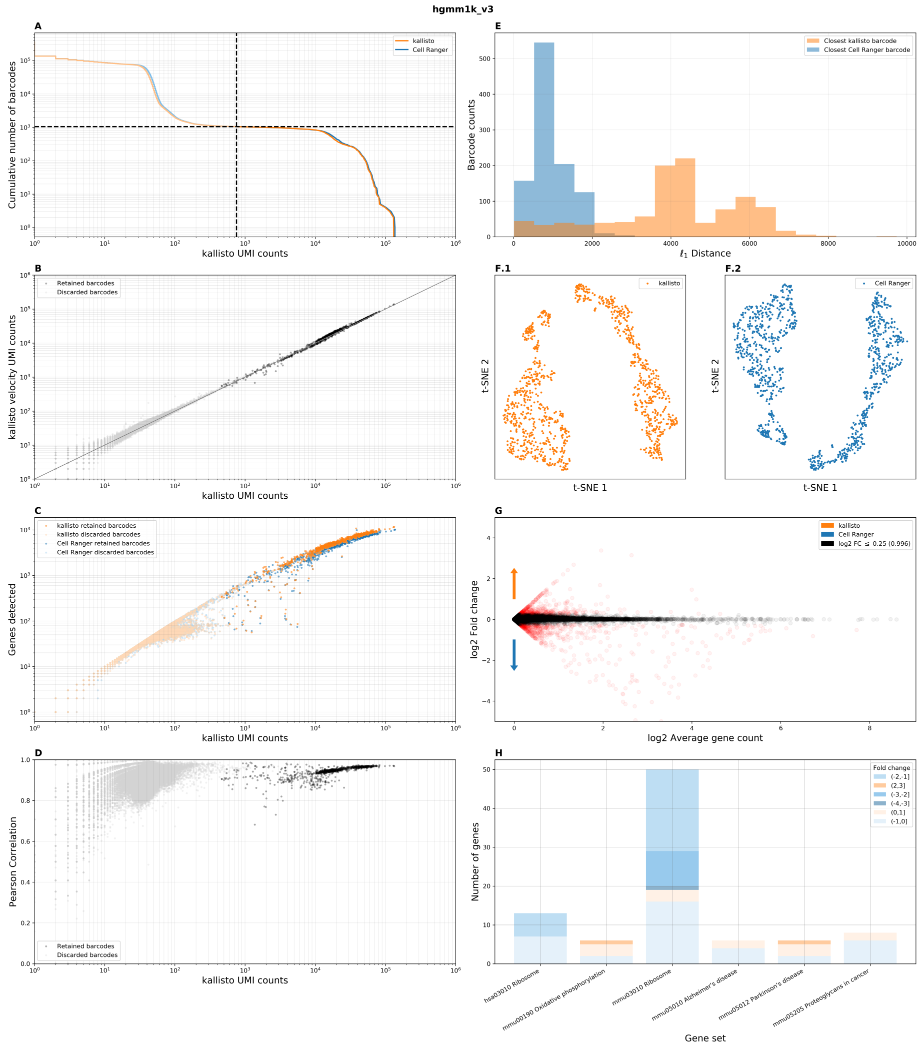


Benchmark panel of dataset `pbmc1k_v3` from 10x Genomics.

### Supplementary Figure 3.6

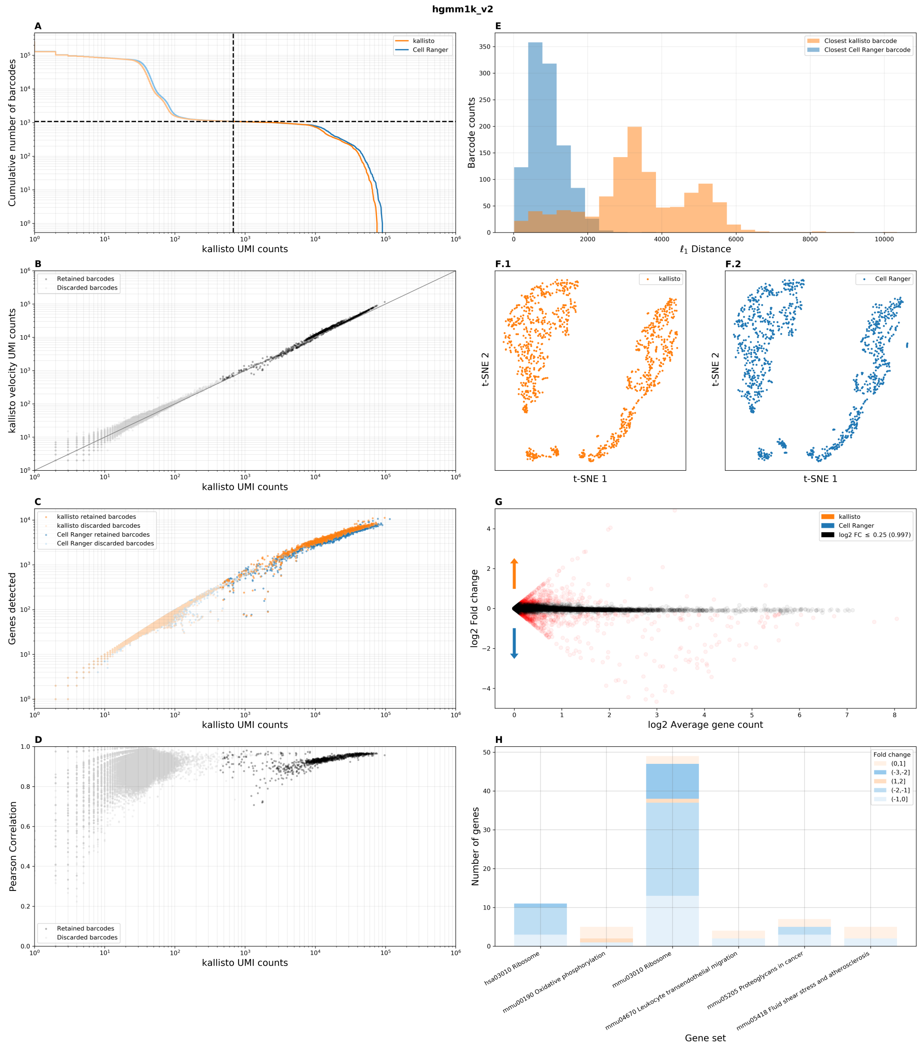


Benchmark panel of dataset `hgmm1k_v2` from 10x Genomics.

### Supplementary Figure 3.7

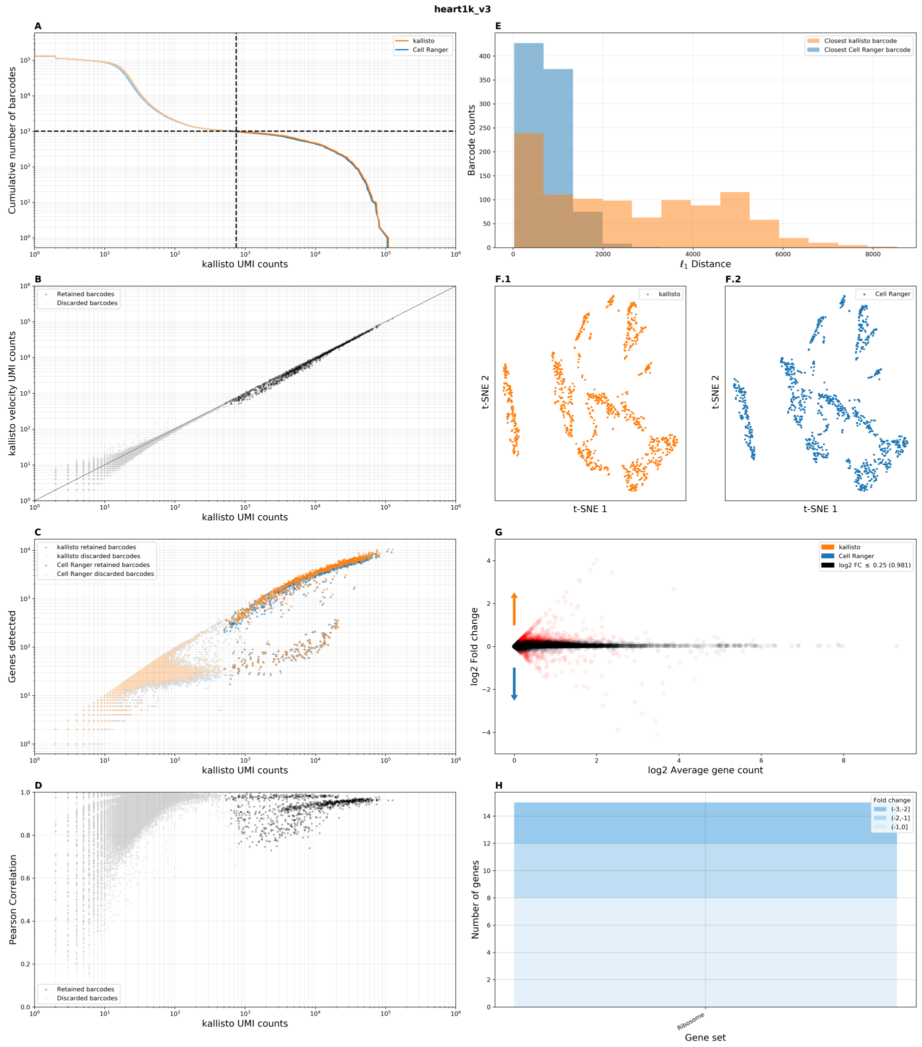


Benchmark panel of dataset `heart1k_v3` from 10x Genomics.

### Supplementary Figure 3.8

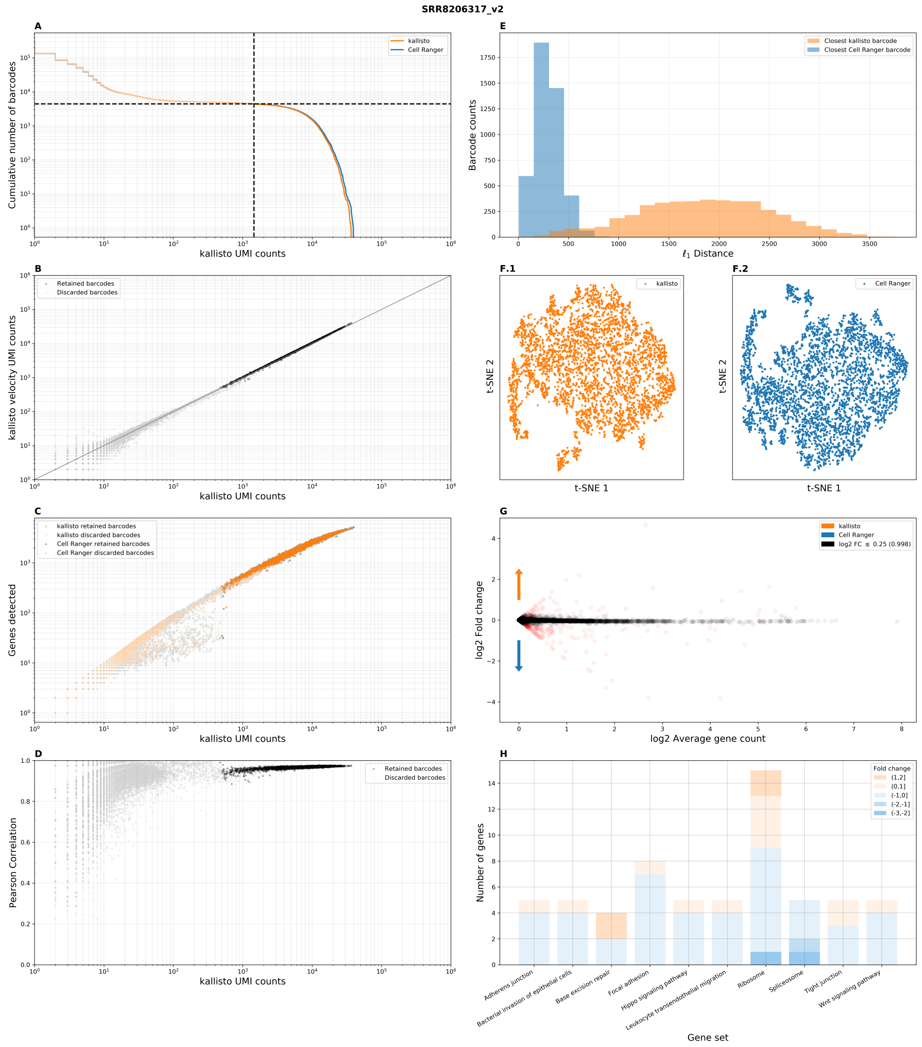


Benchmark panel of dataset SRR8206317 from Miller *et al.* 2019.

### Supplementary Figure 3.9

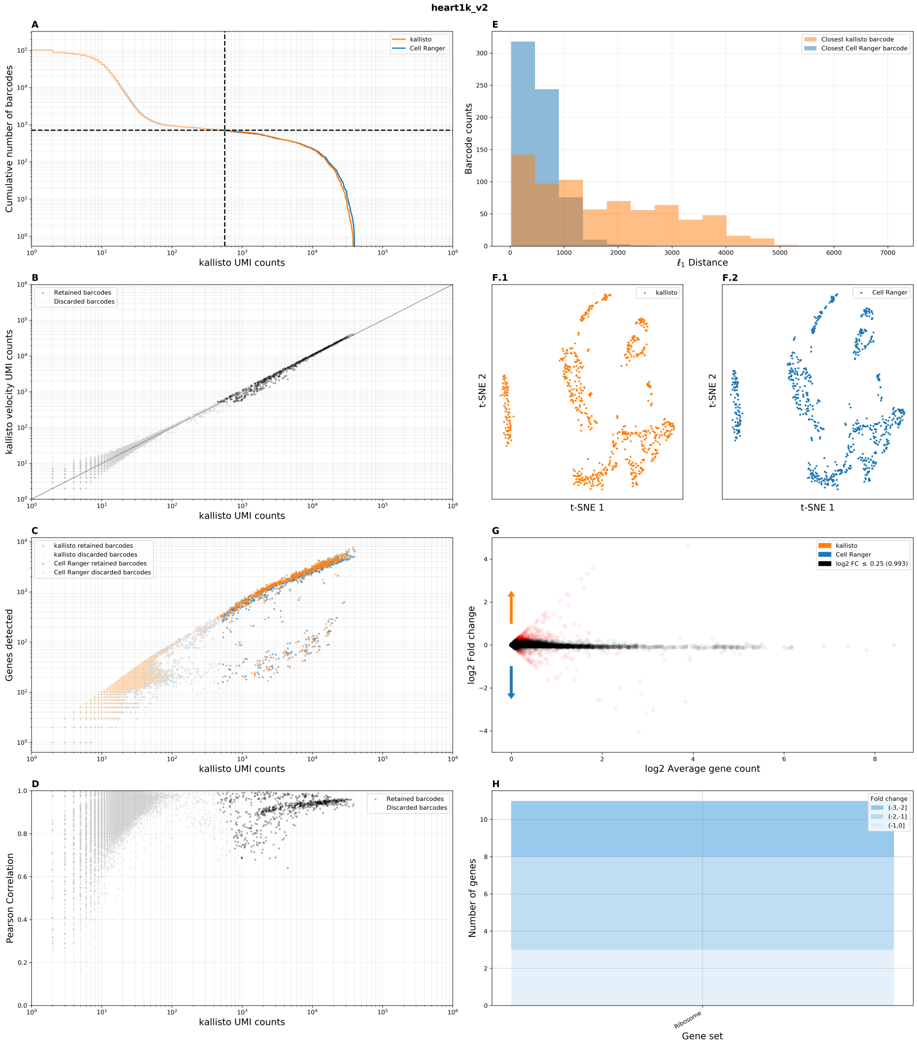


Benchmark panel of dataset `heart1k_v2` from 10x Genomics.

#### Supplementary Figure 3.10

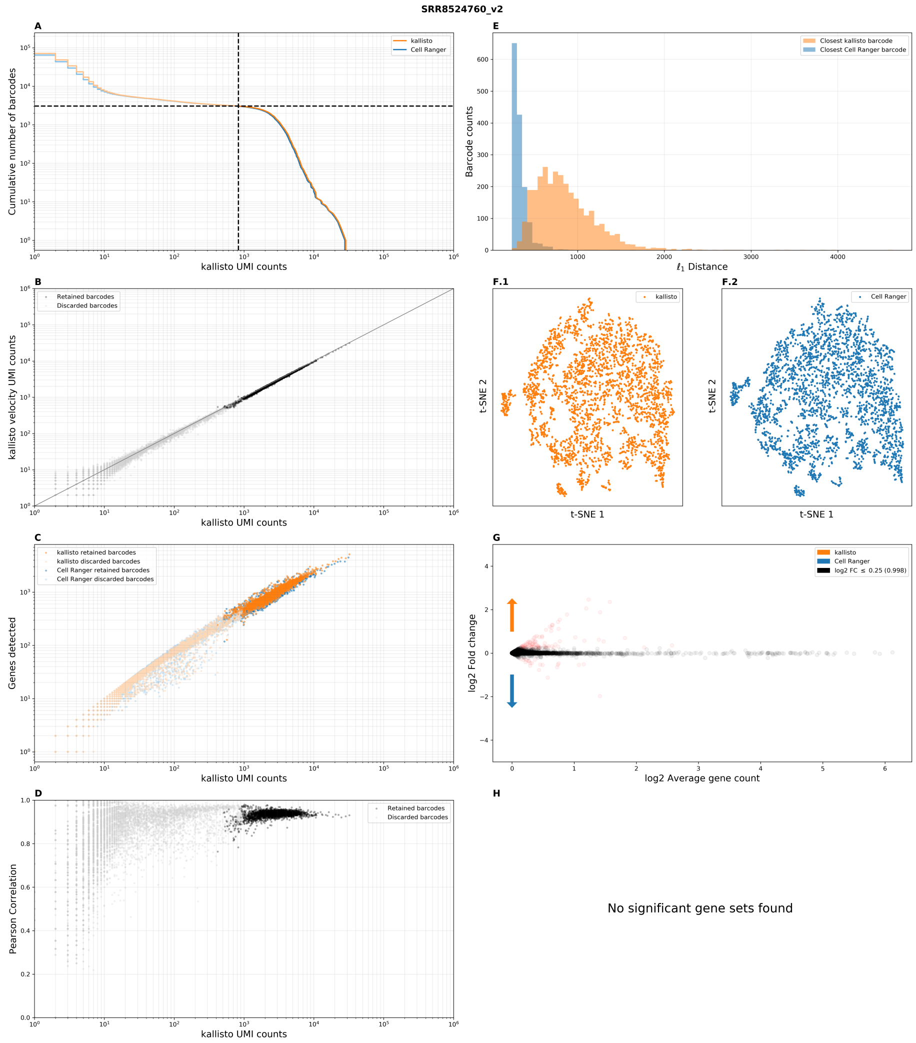


Benchmark panel of dataset SRR8524760 from Carosso *et al.* 2018.

### Supplementary Figure 3.11

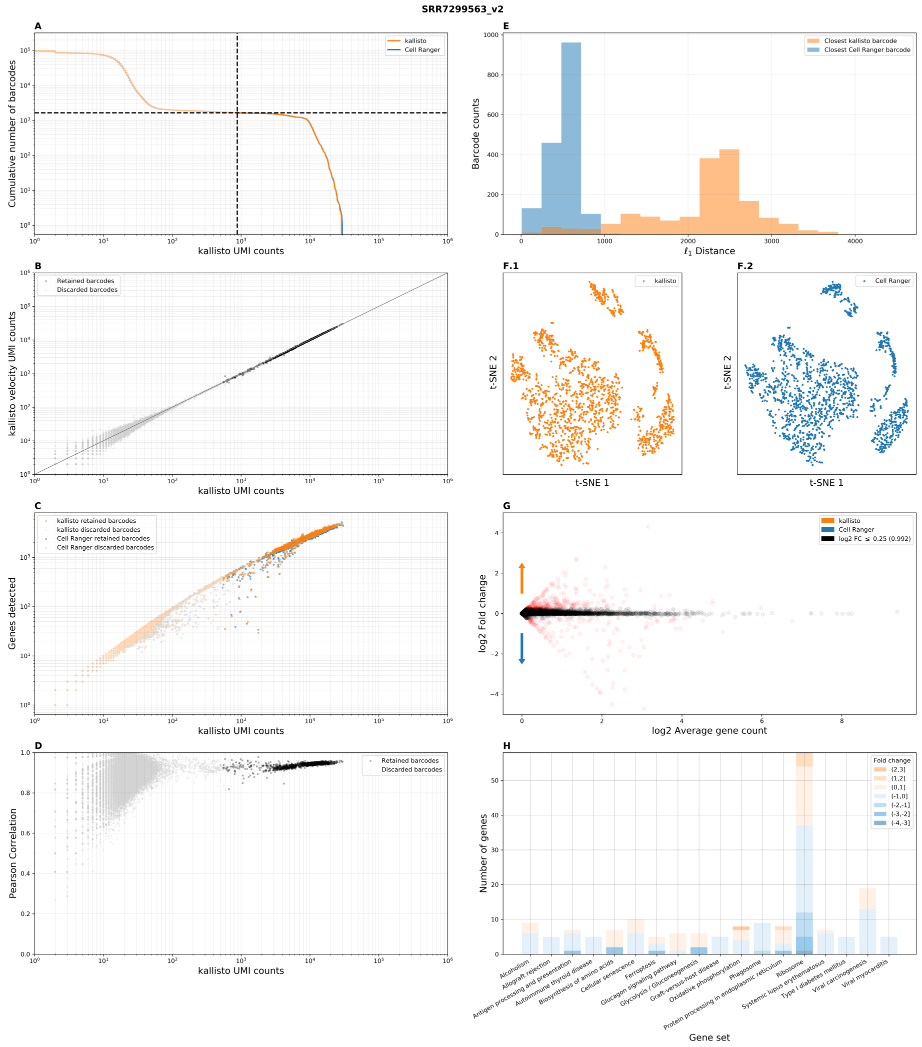


Benchmark panel of dataset `SRR7299563` from Mays *et al.* 2018.

### Supplementary Figure 3.12

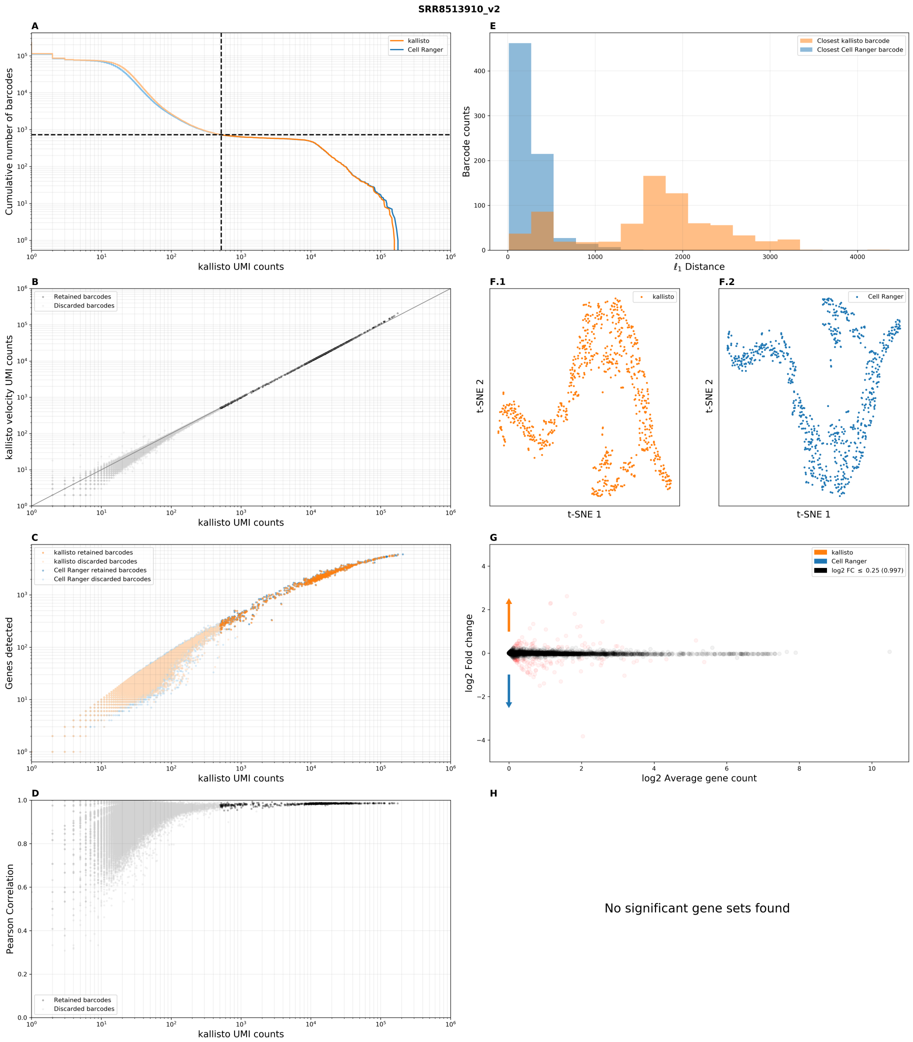


Benchmark panel of dataset `SRR8513910` from Mahadevaraju *et al.* 2019.

### Supplementary Figure 3.13

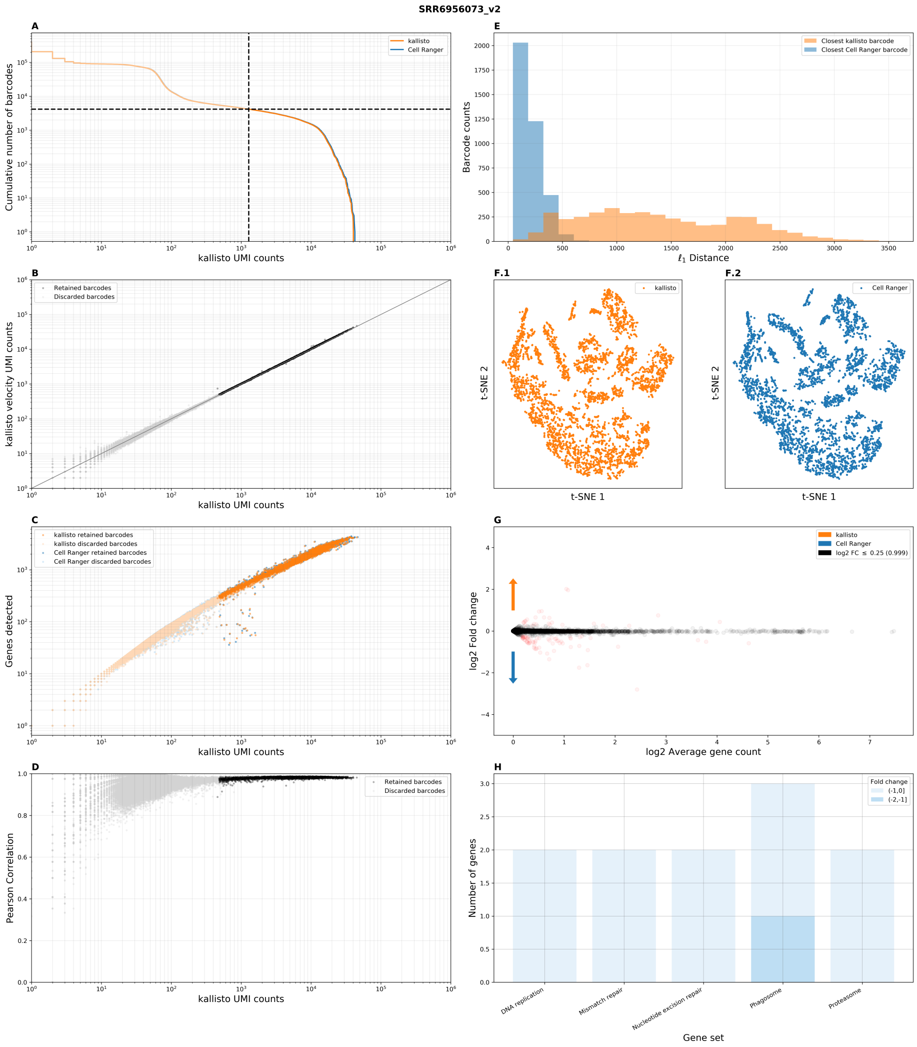


Benchmark panel of dataset `SRR6956073` from Farrell *et al.* 2018.

### Supplementary Figure 3.14

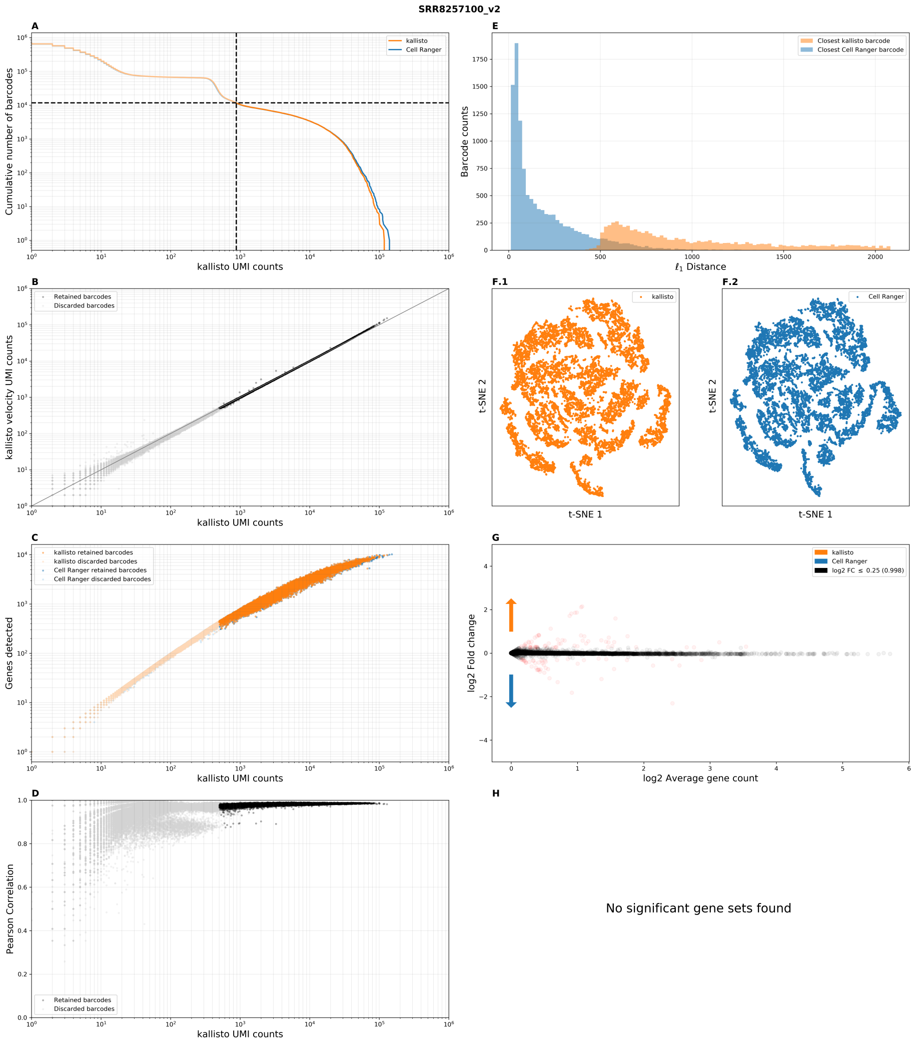


Benchmark panel of dataset `SRR8257100` from Ryu *et al.* 2019.

### Supplementary Figure 3.15

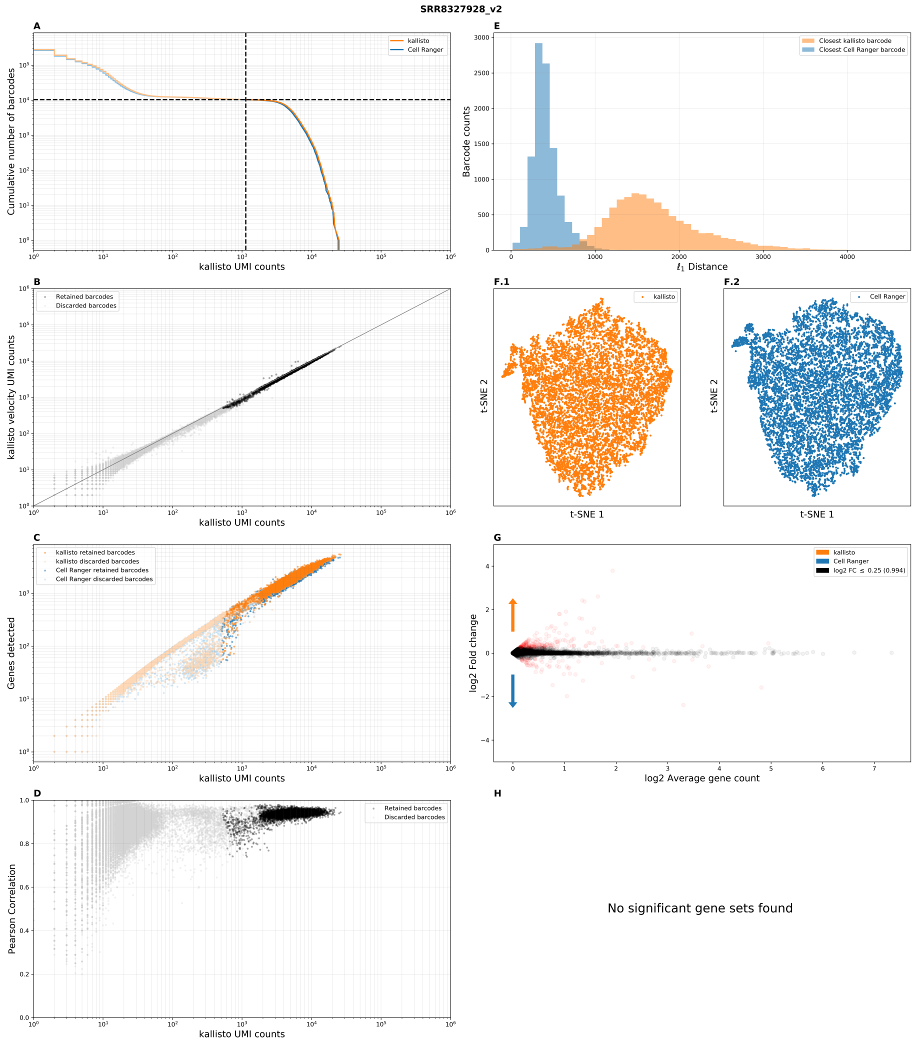


Benchmark panel of dataset `SRR8327928` from Merino *et al.* 2019.

### Supplementary Figure 3.16

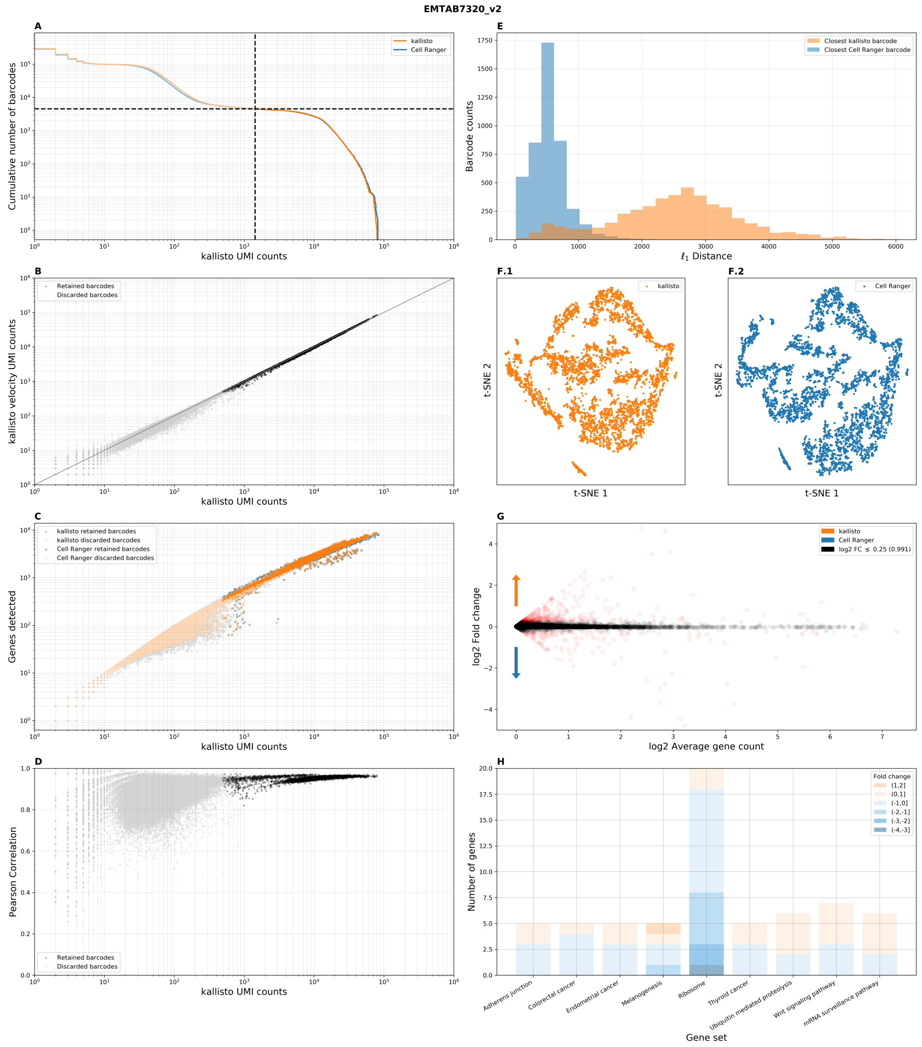

Benchmark panel of dataset `EMTAB7320` from Delile *et al.* 2019.

### Supplementary Figure 3.17

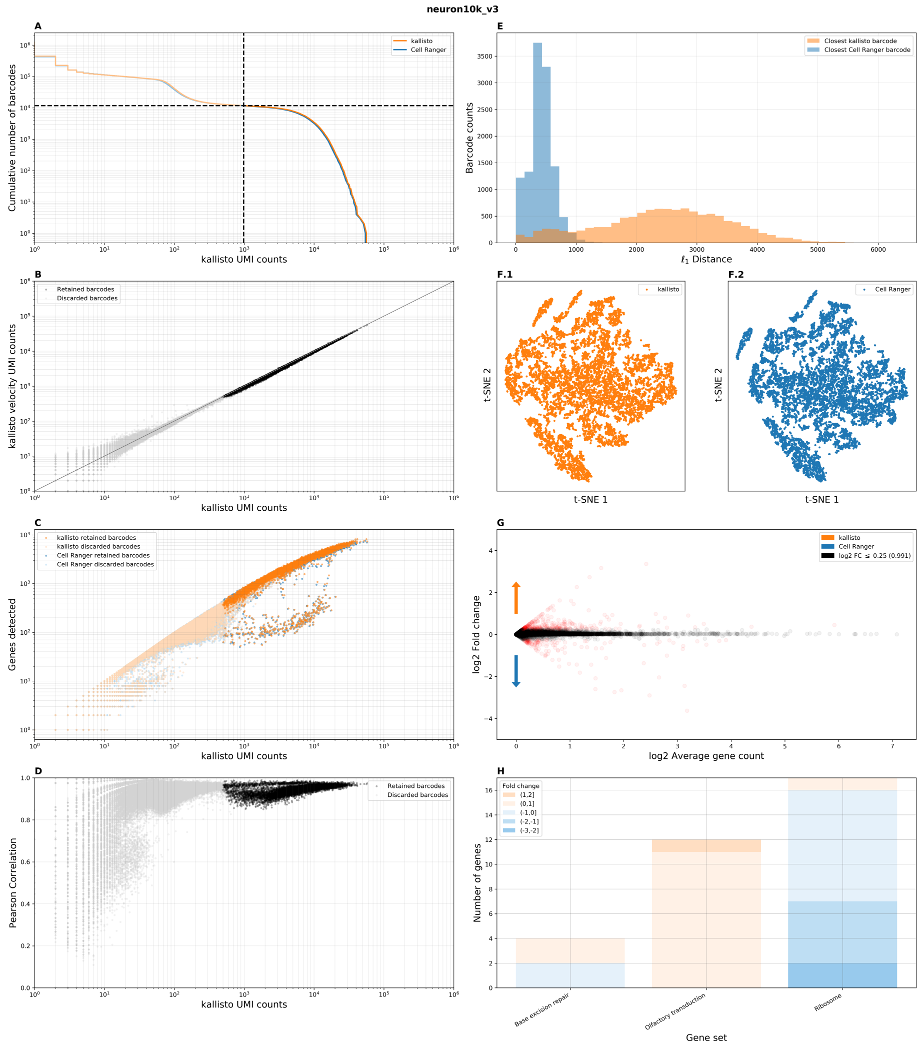


Benchmark panel of dataset `neuron10k_v3` from 10x Genomics.

### Supplementary Figure 3.18

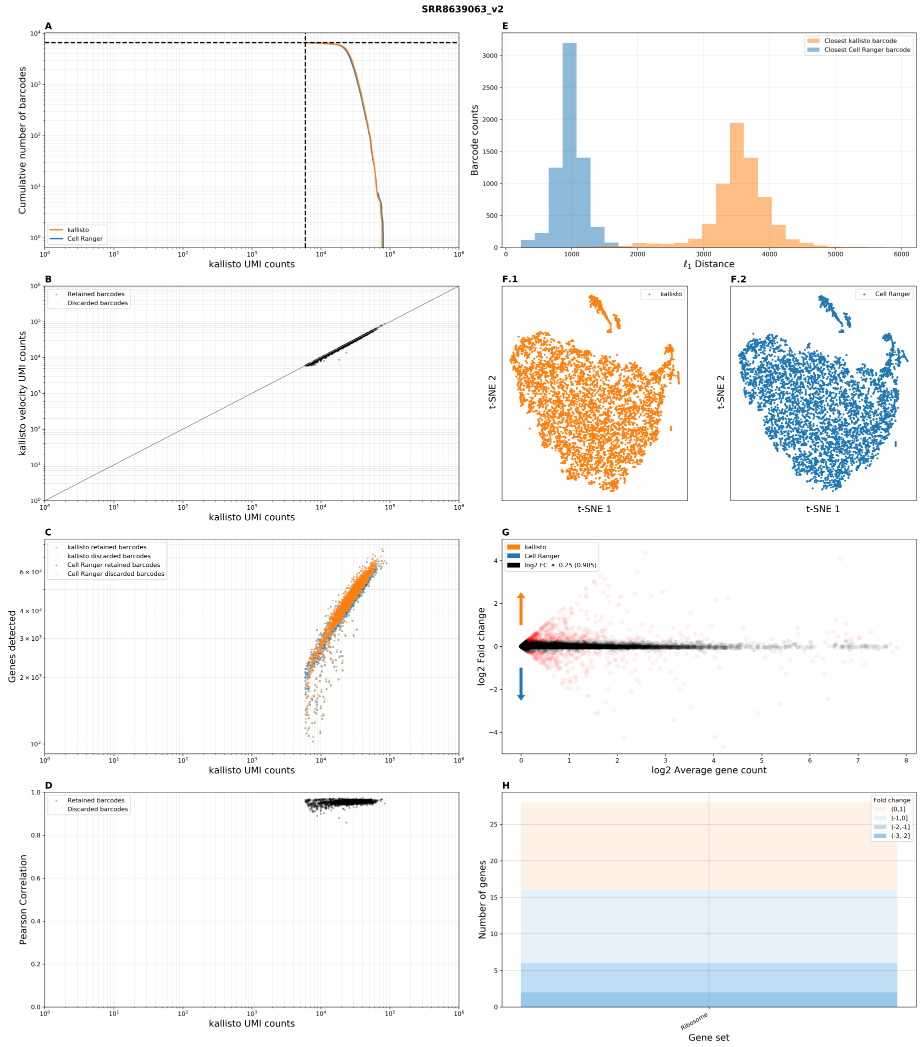


Benchmark panel of dataset `SRR8639063` from Guo *et al.* 2019. Distributed for this dataset were FASTQ files corresponding to filtered barcodes.

### Supplementary Figure 3.19

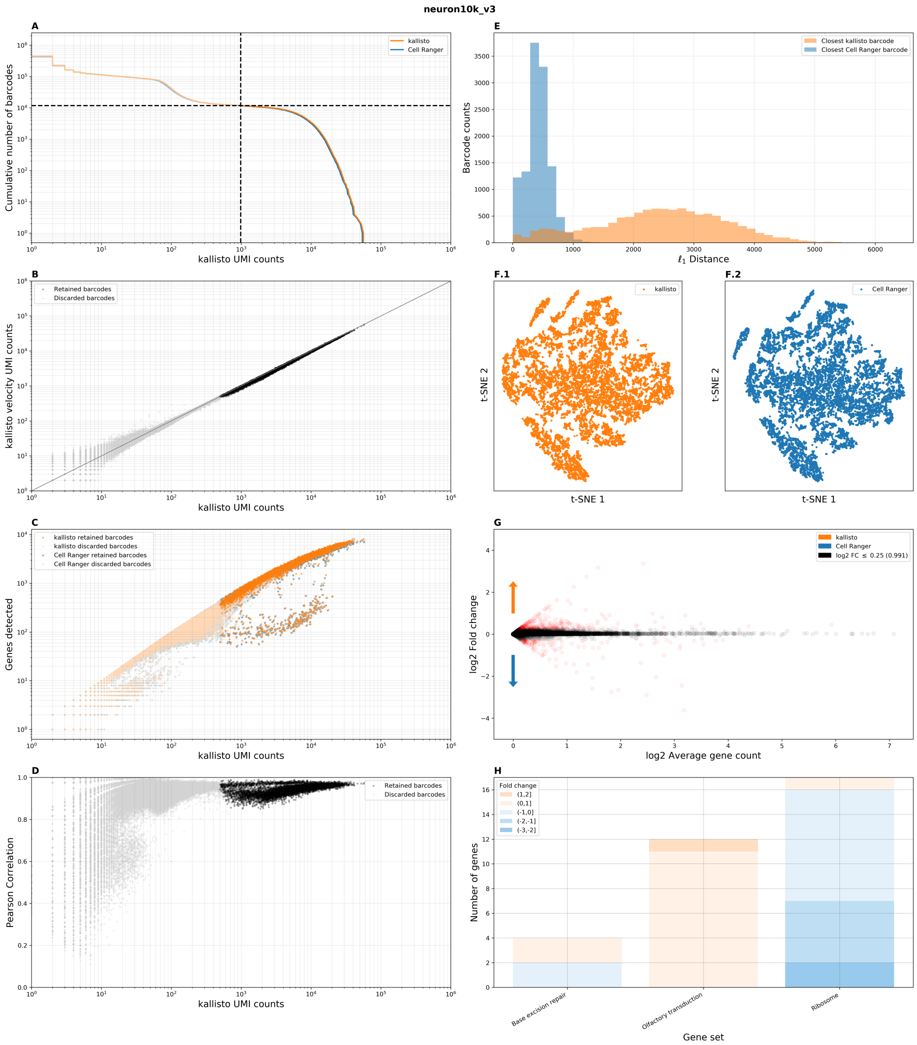


Benchmark panel of dataset `pbmc10k_v3` from 10x Genomics.

### Supplementary Figure 3.20

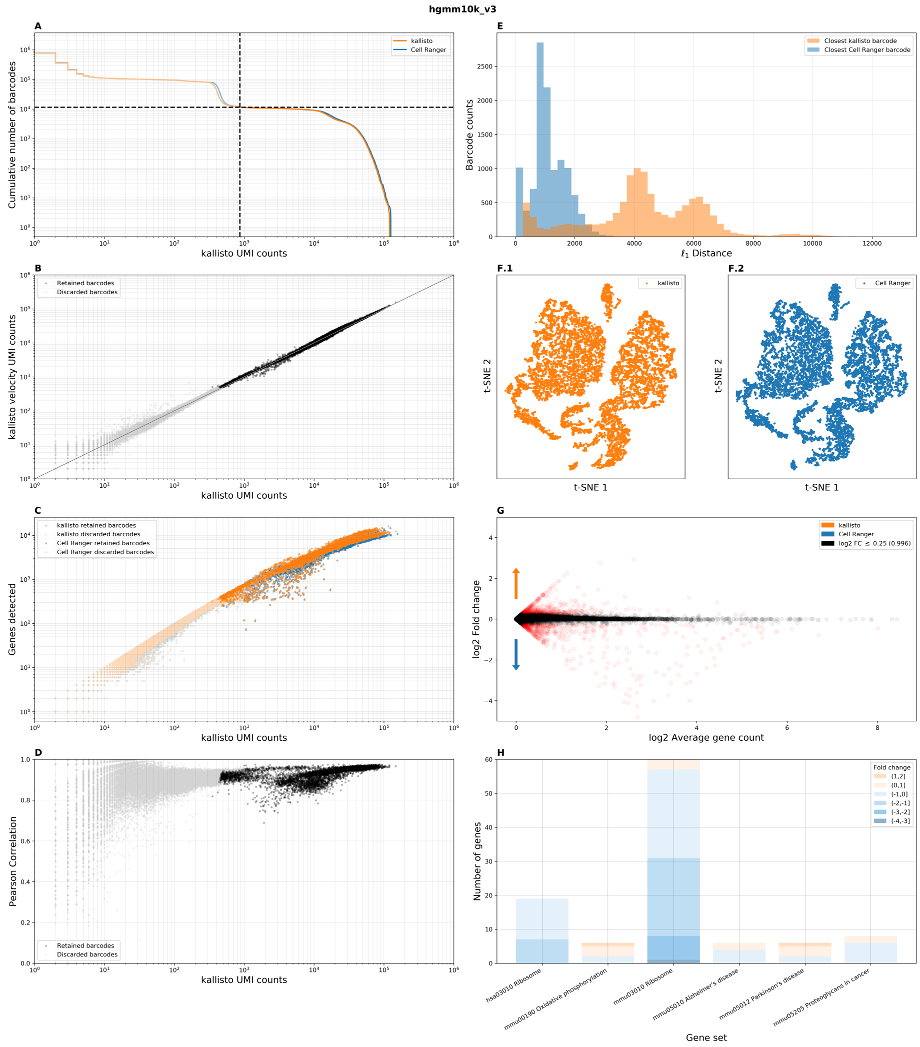


Benchmark panel of dataset `hgmm10k_v3` from 10x Genomics.

#### Supplementary Figure 4

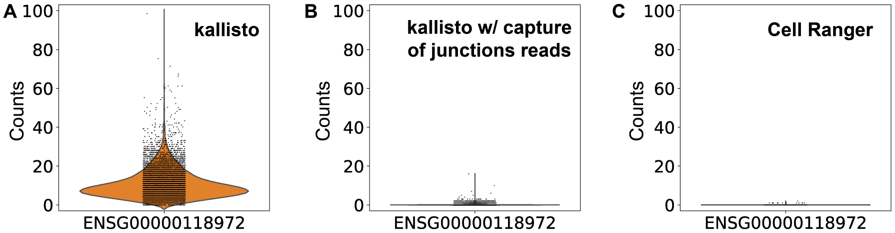


Violin plots displaying distribution of counts for gene FGF23 (ENSG00000118972) in all cells in the dataset `pbmc_10k_v3` using different alignment methods. (A) Transcriptome pseudoalignment with kallisto using a standard index constructed from ENSEMBL transcripts. (B) Transcriptome pseudoalignment with kallisto using a modified index that includes, separately, sequences from splice junctions to capture unspliced junction reads. (C) Genome alignment with Cell Ranger. The gene was selected as an example as it was an outlier in discrepancy between kallisto and Cell Ranger when quantification was done with the standard index.

#### Supplementary Figure 5

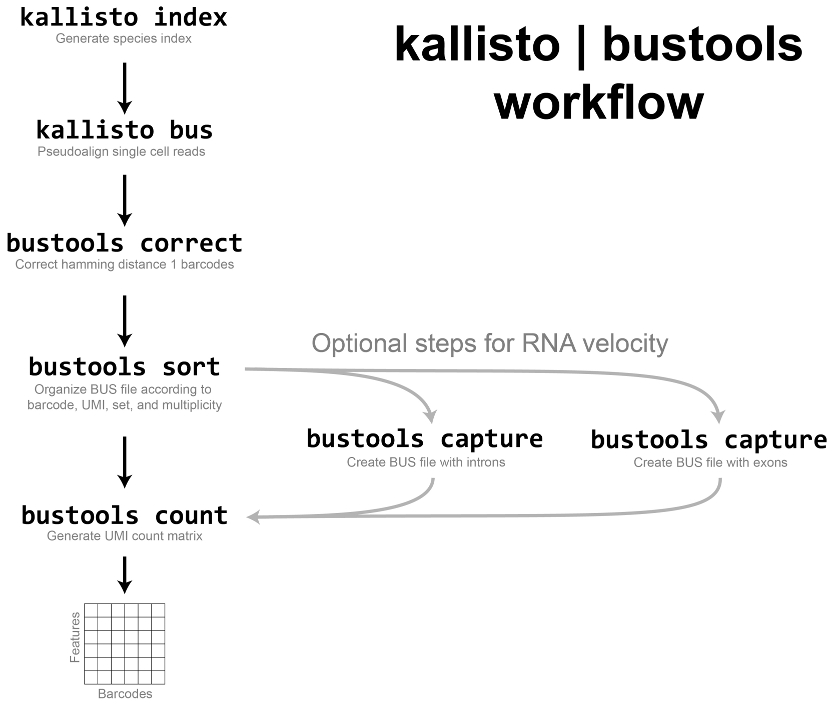


Overview of the kallisto bustools workflow. First an index for kallisto is built from a set of transcript sequences using the kallisto index command. Then kallisto bus is run on the FASTQ files; this generates a BUS file that contains records corresponding to reads, with data on the cell barcode, UMI, and transcript compatibility of each read. The barcodes are then corrected by processing the BUS file with the bustools correct command, after which the BUS file is sorted with bustools sort. Here, duplicate reads (those reads sharing an identical cell barcode, UMI, and equivalence class triplet) are collapsed into a single record and their abundance saved as a new metadata column in the BUS file named "multiplicity". Finally, bustoools count produces cells x features count matrices. If kallisto bus is run with an index containing intron sequences, the bustools capture command can be used to produce spliced and unspliced matrices for RNA velocity after sorting and before counting.

### Supplementary Figure 6

#### Supplementary Figure 6.1

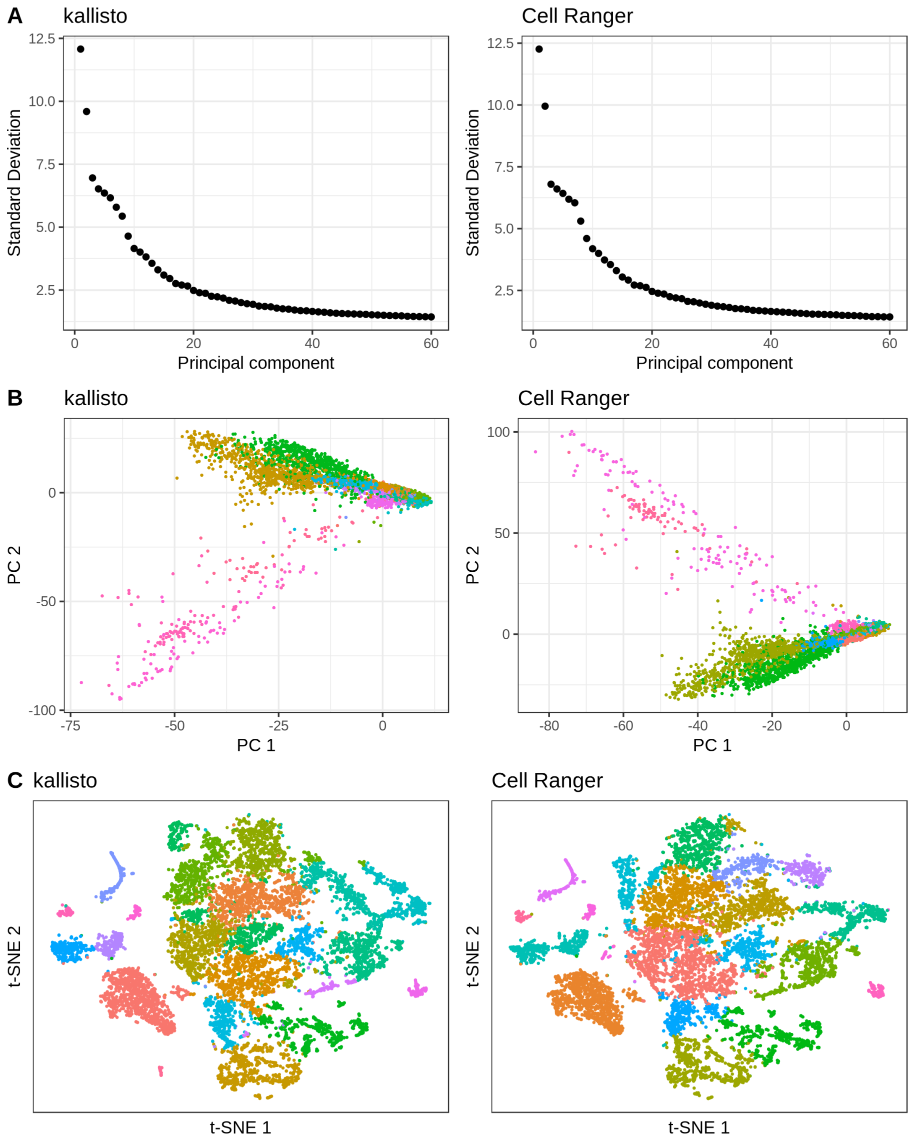


(A) Elbow plot of standard deviation explained by each principal component of the gene count matrix from kallisto and Cell Ranger. (B) Cell embedding in the first 2 principal components colored by cluster. (C) Cell embedding in tSNE colored by cluster.

#### Supplementary Figure 6.2

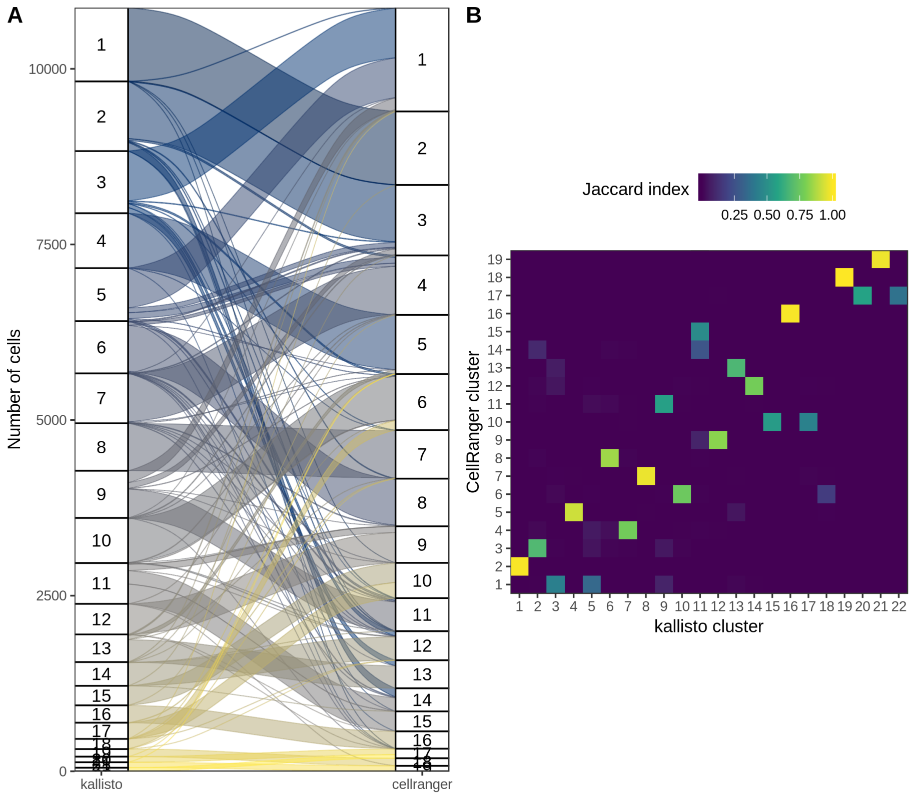


(A) Number of cells assigned to each cluster by kallisto and Cell Ranger and the correspondence between the clusters. (B) Jaccard indices between each kallisto cluster and each Cell Ranger cluster.

### Supplementary Figure 6.3

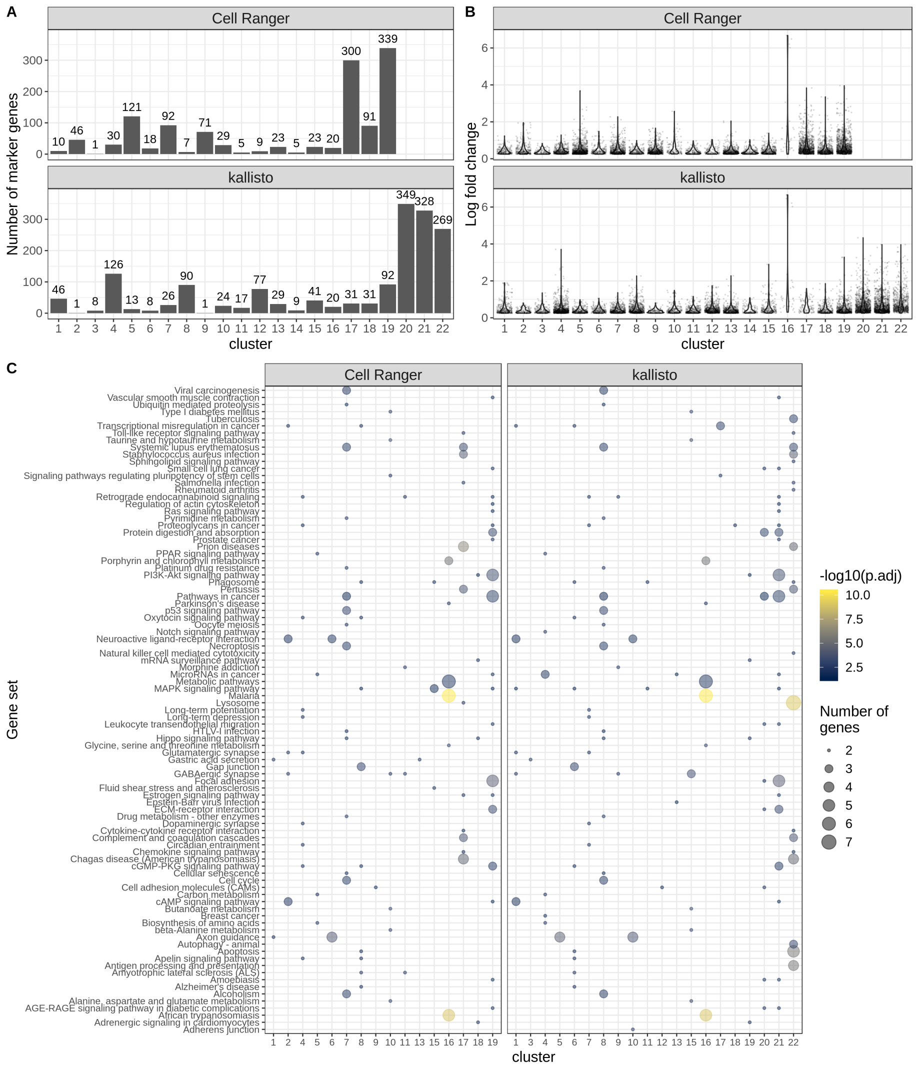


(A) Number of marker genes with log fold change of at least 0.75 and adjusted $p < 0.05$ in each cluster. (B) Log fold change of marker genes in each cluster. (C) Gene set enrichment of top 20 marker genes (log fold change) in each cluster.

#### Supplementary Figure 6.4

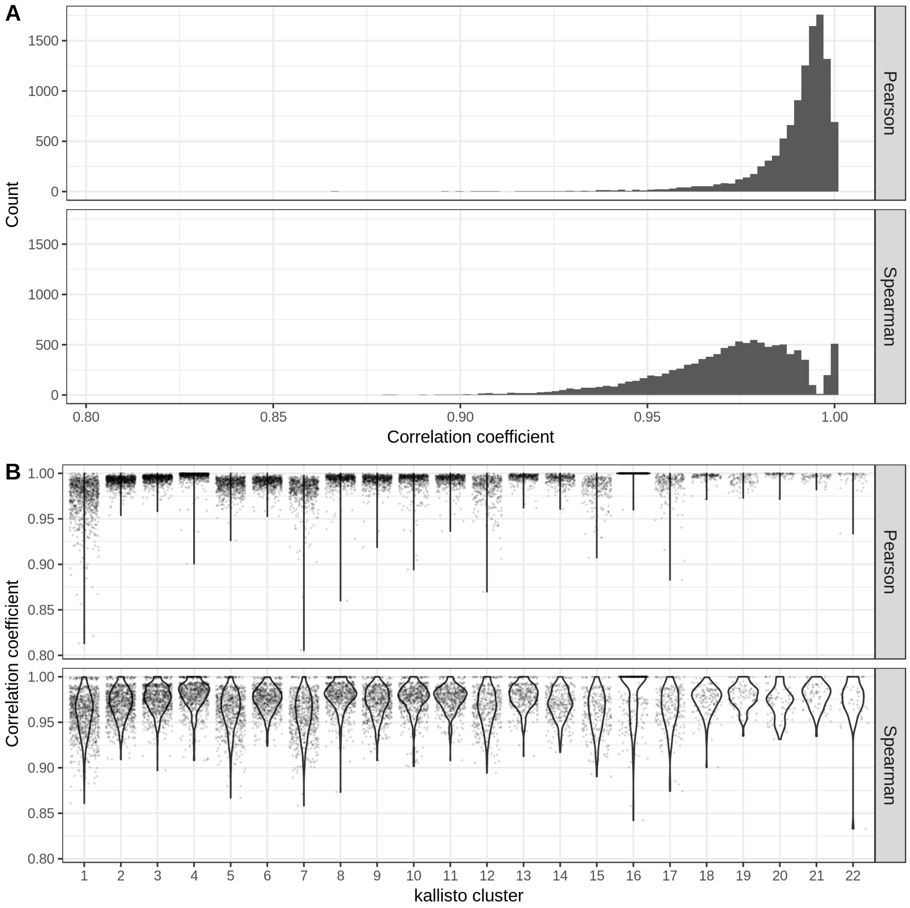


(A) Histograms of Spearman and Pearson correlation coefficients between barcodes from kallisto and the same barcodes from Cell Ranger for the top 15 marker genes (by log fold change) of each cluster. (B) Spearman and Pearson correlation coefficients, as in (A), for cells in each kallisto cluster. cluster 16 corresponds to erythrocytes, while most other cells are neuronal precursor cells.

#### Supplementary Figure 6.5

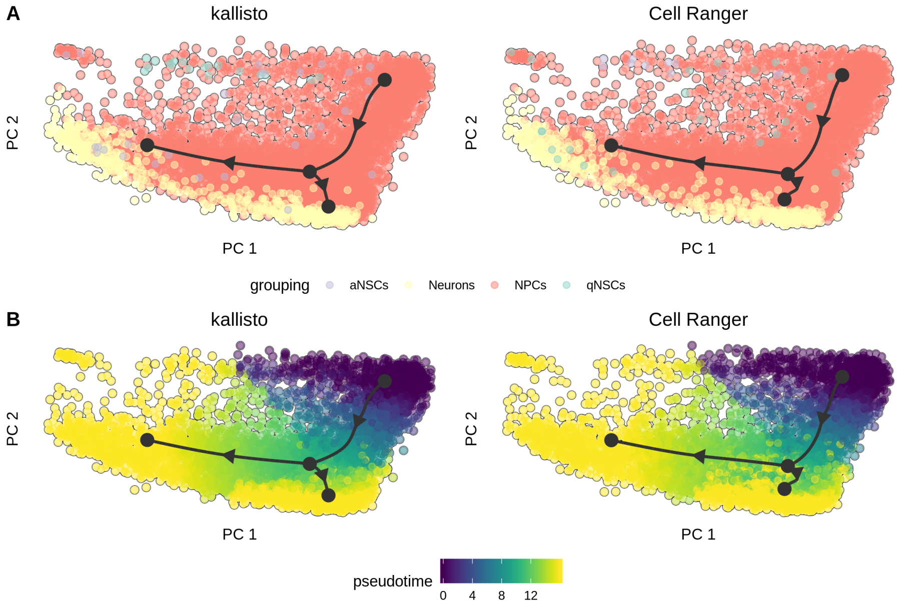


(A) Lineage inference with |slingshot— projected to the first 2 principal components, with cells colored by cell type inferred by |SingleR— Aran *et al.* 2019. The reference used for |SingleR— is from Benayoun *et al.* 2019. aNSCs stands for active neuronal stem cells. NPCs stands for neuronal precursor cells. qNSCs stands for quiescent neuronal stem cells. (B) Coloring by pseudotime values from slingshot.

#### Supplementary Figure 7

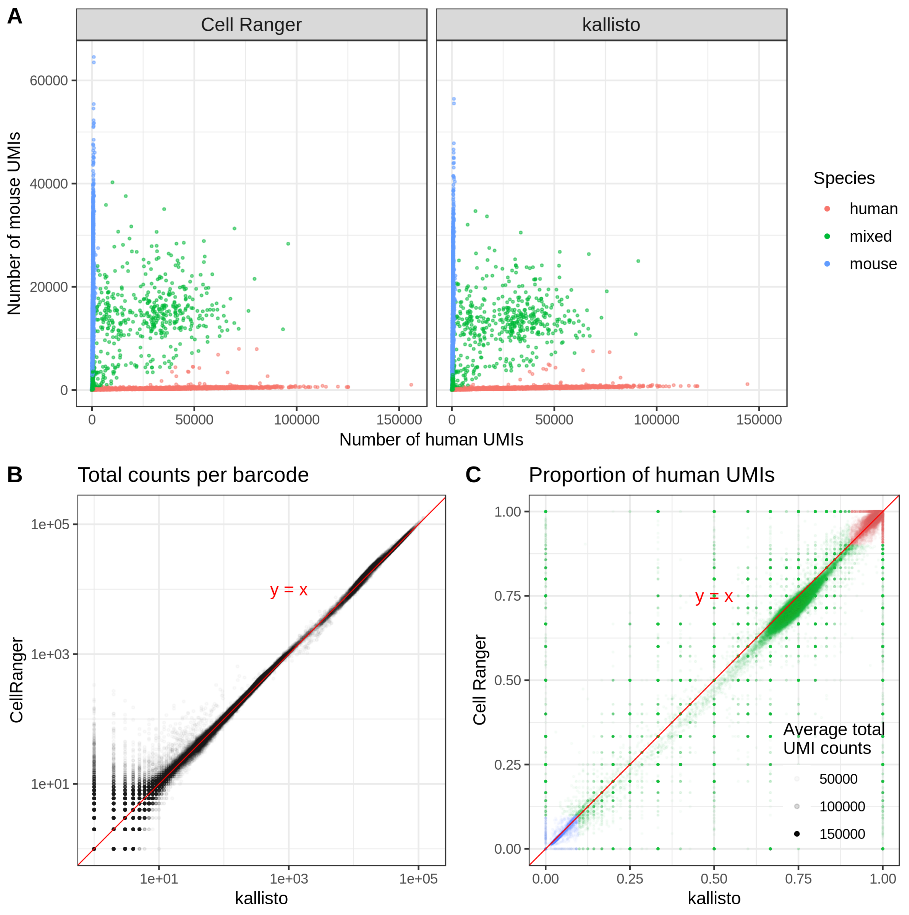


Comparison of Cell Ranger to kallisto on the 10x Genomics `hgmm10k_v3` species mixing experiment, 10k 1:1 Mixture of Fresh Frozen Human (HEK293T) and Mouse (NIH3T3) Cells. `https://support.10xgenomics.com/single-cell-gene-expression/datasets/3.0.0/hgmm_10k_v3` (A) Barnyard plot with droplets colored according to species of origin: human (red), mouse (blue) and mixed (green). Mixed droplets correspond to cell doublets. (B) The number of total counts per barcode in Cell Ranger and kallisto. (C) The proportion of UMIs in each droplet originating from human. The cluster of droplets in the lower left corner correspond to mouse cells and that in the upper right corner to human cells. The middle band of droplets are doublets. Droplets are shaded according to the number of distinct UMIs they contain.

#### Supplementary Figure 8

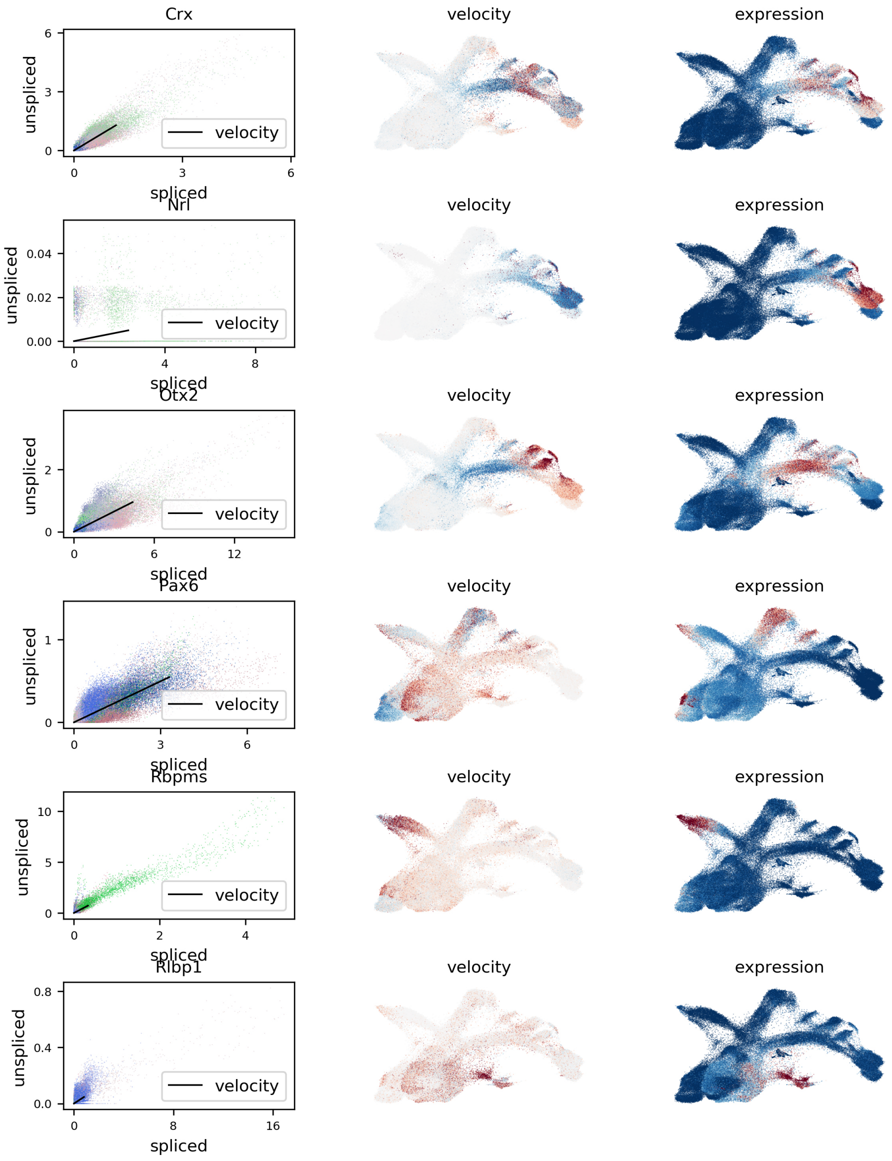


Phase diagrams and expression/velocity for six marker genes studied in Clark *et al.* 2019. The expression results are concordant with pseudotime analysis.

### Supplementary Figure 9

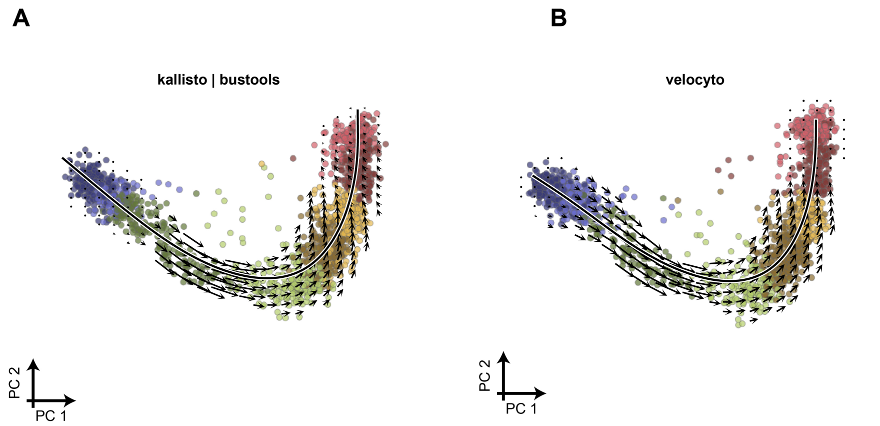


RNA velocity based on spliced and unspliced matrices from a dataset of 1,720 developing human glutamatergic neurons at post-conception week 10 from La Manno *et al.* 2018. The colors correspond to cell types and intermediate states and a principal "velocity curve" is shown in bold. (A) RNA velocity based on spliced and unspliced matrices computed with kallisto and bustools. (B) RNA velocity based on the spliced and unspliced matrices computed with velocyto.

### Supplementary Figure 10

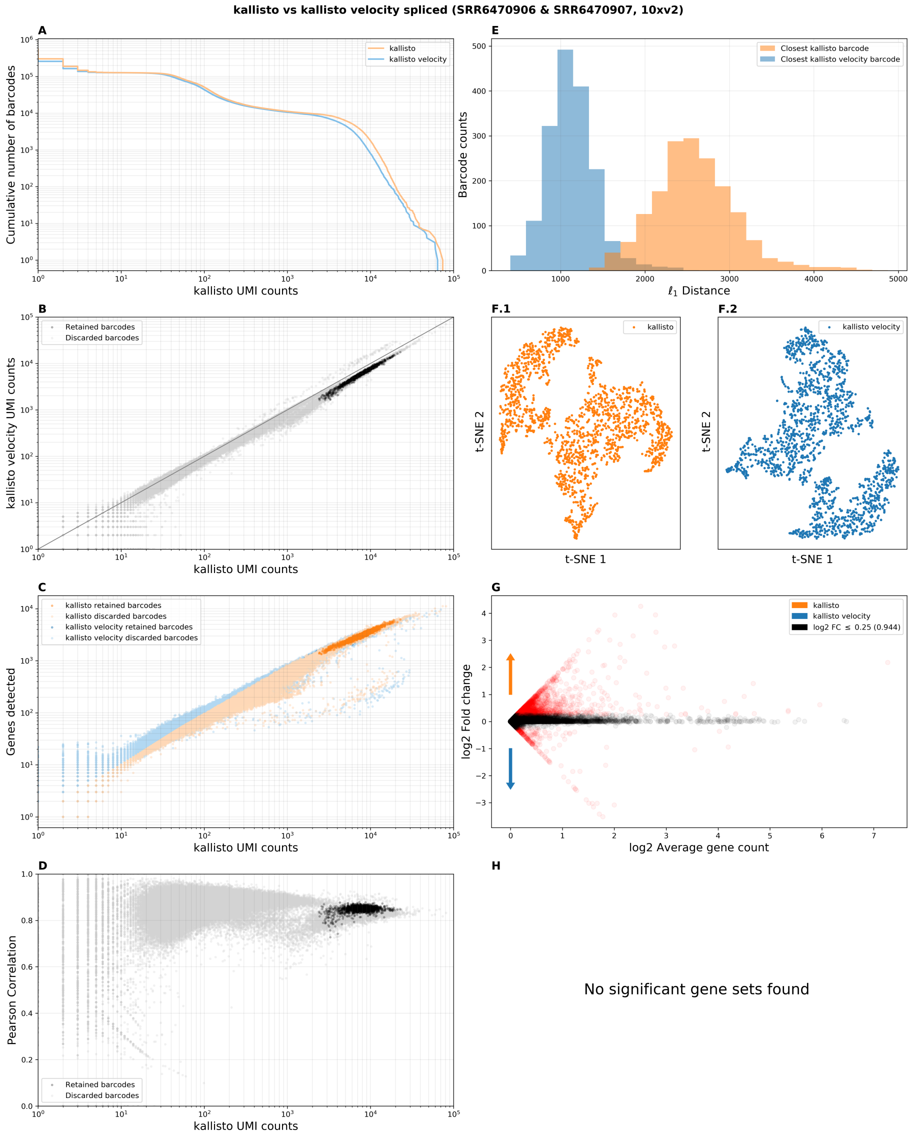


Comparison of count matrix generated by the standard kallisto workflow with the spliced count matrix generated by the kallisto RNA velocity workflow.

### Supplementary Figure 11

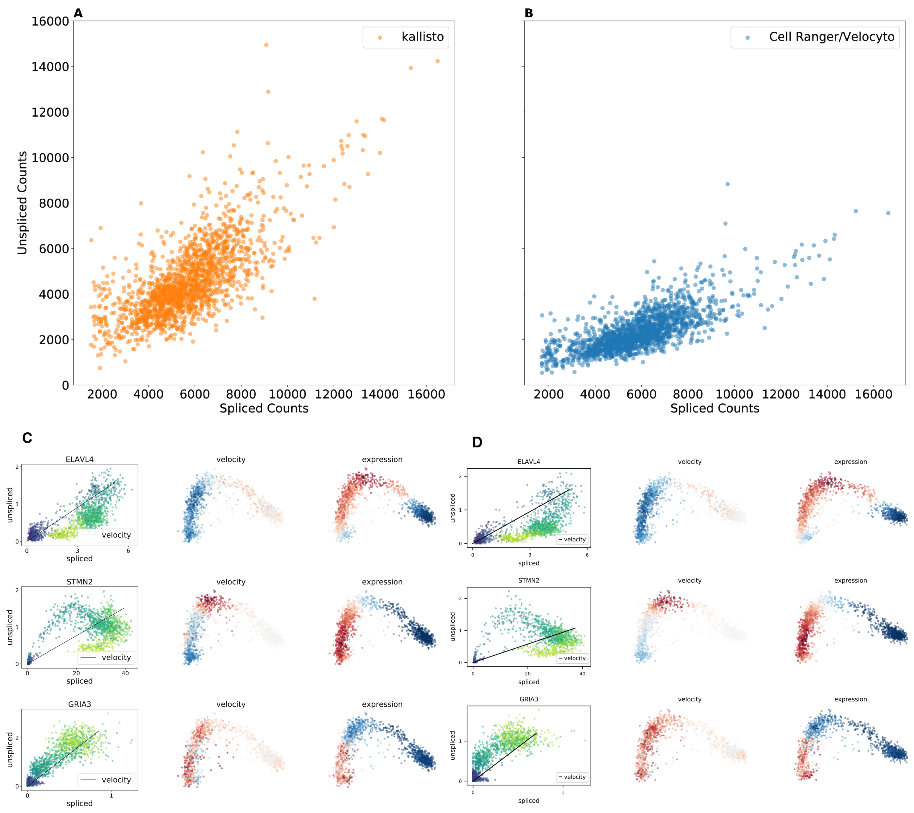


Comparison of Cell Ranger and velocyto to kallisto in an RNA velocity analysis of developing human glutamatergic neurons at post-conception week 10. (A) Number of distinct UMIs from spliced vs. unspliced transcripts from kallisto. (B) Number of distinct UMIs from spliced vs. unspliced transcripts from Cell Ranger/Velocyto. Cell Ranger/Velocyto has similar numbers of spliced counts but fewer unspliced counts. (C) Phase diagrams from the kallisto RNA velocity analysis for 3 genes highlighted in La Manno *et al.* 2018. (D) Corresponding phase diagrams from the Cell Ranger/Velocyto RNA velocity analysis showing agreement with the kallisto results.

#### Supplementary Figure 12

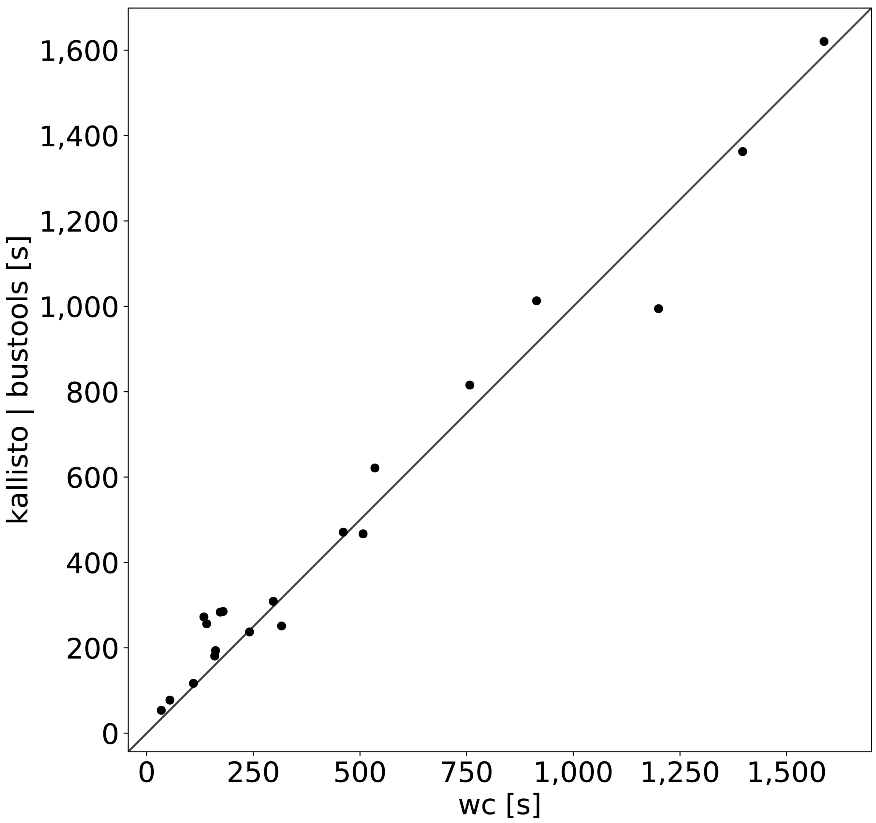


Comparison of kallisto runtimes with those of the Unix word count (wc) command. Each point corresponds to a different dataset.

#### Supplementary Figure 13

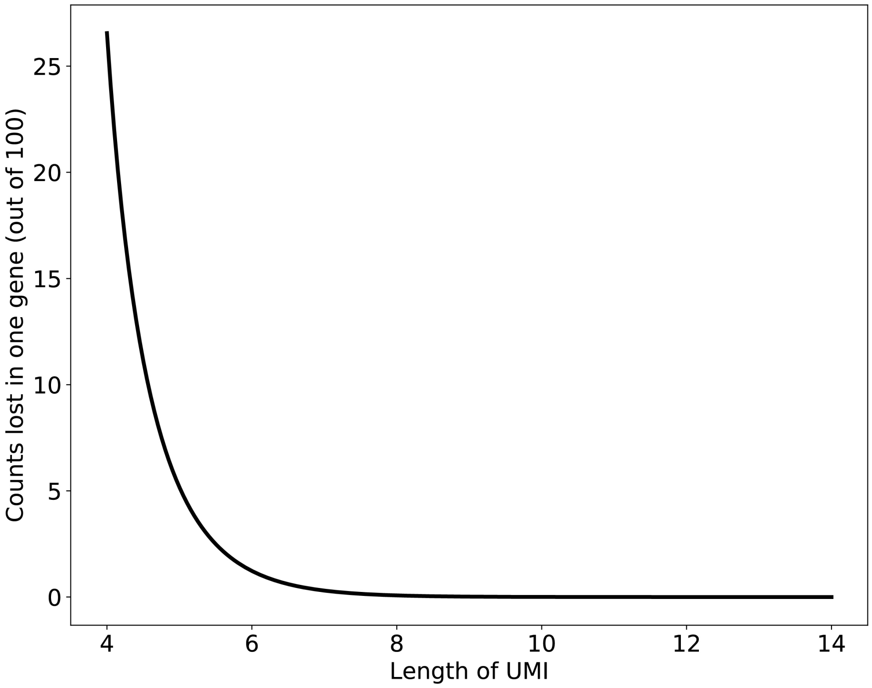


The number of counts lost due to naïve collapsing of UMIs as a function of the length of the UMIs for a gene with 100 counts. The calculation, based on Supplementary Note equation (11), assumes that the effective number of UMIs is $4^L$ when UMIs are of length $L$.
