## Supplementary Note for "Modular and efficient pre-processing of single-cell RNA-seq"

### 1 Preliminaries

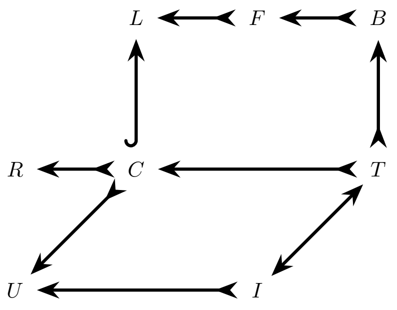


Figure 1: Diagram of sets associated with a cell in a single-cell RNA-seq sequencing experiment.

A single-cell RNA-seq experiment can be described as follows: the goal of the experiment is to identify the ensemble of RNA molecules in multiple cells; in Figure 1 the ensemble of RNA molecules contained within a single cell is denoted by $R$. To investigate $R$ a library ($L$) is constructed from the set of molecules captured from $R$ (the set $C$). Typically, $L$ is the result of of various fragmentation and amplification steps performed on $C$, meaning each element of $C$ may be observed in $L$ with some multiplicity. Thus, there is an inclusion map from $C$ to $L$, and an injection from $C$ to $R$. The library is interrogated via sequencing of some of the molecules in $L$, resulting in a set $F$ of fragments. Subsequently, the set $F$ is aligned or pseudoaligned to create a set $B$, which in this paper is a BUS file (Melsted, Ntranos, and Pachter 2019). Not every fragment $F$ is represented in $B$, hence the injection, rather than bijection, from $B$ to $F$, and similarly from $F$ to $L$. The set $T$ consists of transcripts that correspond to molecules in $C$ that were represented in $B$. Note that $|R| \geq |C| \geq |T|$. Separately, the set $U$ consists of the UMIs on the bead that the cell was trapped with, and $I$ is a multiset of UMIs associated with the molecules in

$R :=$ the multiset of all RNAs in a cell.
$L :=$ the multiset of all molecules in the library.
$F :=$ the multiset of reads.
$B :=$ the multiset of {barcode, UMI, equivalence class} triplets.
$C :=$ the multiset of captured RNA molecules represented in the library.
$T :=$ the set of captured transcripts represented in B.
$U :=$ the set of UMIs in a droplet.
$I :=$ the set of UMIs represented in B.

Table 1: Notation for description of a single-cell experiment.

$T$ (Table 1).

The data in a single-cell experiment consists of the sets $F$ for each cell. In our workflow, a combined BUS file (merge of the sets $B$) is generated using kallisto (Bray et al. 2016). While the multiset $I$ is not directly measured, it's support $supp(I)$ (the set of distinct UMIs) can be extracted from the BUS file. The goal of single-cell RNA-seq pre-processing is to infer the multiset $T$. What we describe in this note is an approach to estimating two different quantities: the effective sizes $|U|$ of the sets of UMIs associated with each bead, and the number of captured molecules represented in the BUS file, i.e. $|I|$ or equivalently $|T|$. Specifically, we are interested in the restriction of the latter to individual genes in cells, for the purpose of estimating the error in the number of counts that can be introduced when naïvely collapsing UMIs by gene. The reason for estimating $|U|$ is that it is necessary to estimate $|I|$.

### 2 Modeling an experiment

The number of distinct UMIs on a bead in a droplet is at most $4^L$ where $L$ is the number of UMI bases (the 10xv2 technology uses $L = 10$ and the 10xv3 technology $L = 12$). For a bead captured along with a cell in a droplet, we denote the number of UMIs on the bead by $n = |U|$. We model the process by which UMIs are associated with molecules as follows: each UMI is selected by sampling uniformly at random from the set of UMIs $U$. In other words, the molecules are labeled with UMIs by sampling with replacement. This model has been used previously (Grün, Kester, and Oudenaarden 2014), and is justified by distributions of UMIs seen empirically (Figure 2). If $k = |I|$ is the number of UMIs in a library derived from a single droplet then the assumption of uniform random sampling of UMIs from the associated bead implies that the probability that a specific UMI is observed zero times is $\left(1 - \frac{1}{n}\right)^k$. Therefore the expected number of UMIs observed at least once, i.e. the expected number of distinct UMIs in a cell, is

$$n\left(1 - \left(1 - \frac{1}{n}\right)^k\right). \tag{1}$$

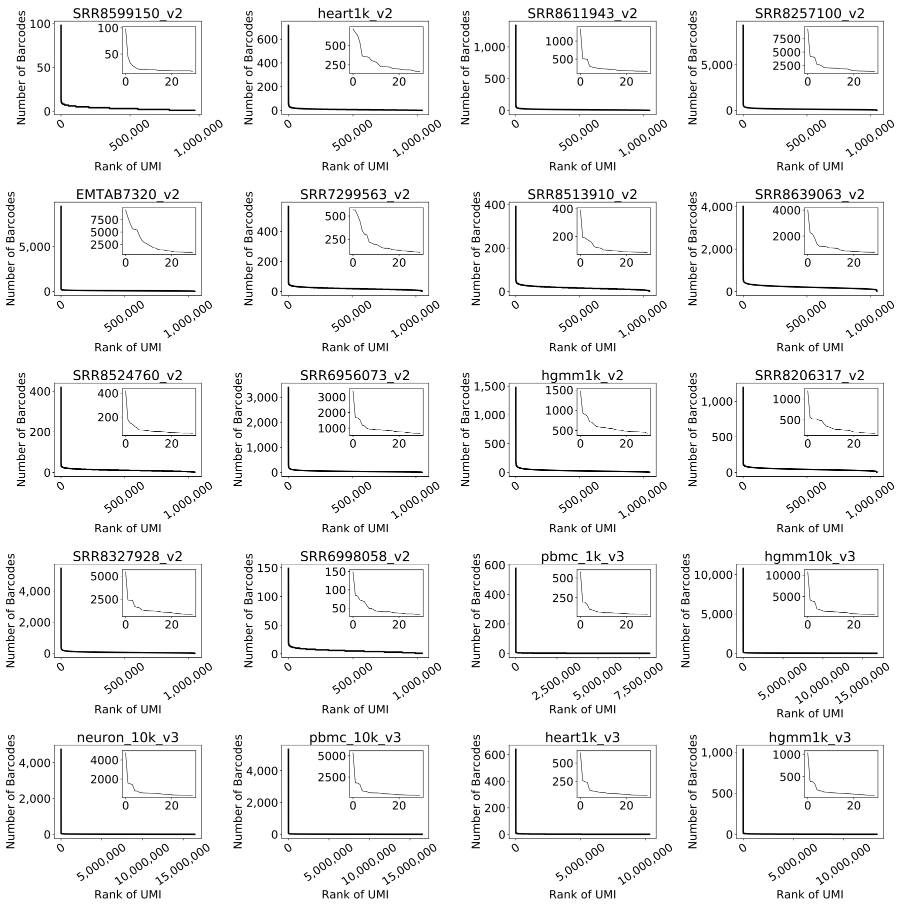


Figure 2: Distribution of UMIs across cells. With the exception of a handful of artifacts, UMIs are uniformly distributed across cells.

### 3 Estimating the effective number of UMIs

To estimate $n$ ($=|U|$) we utilize two observations:

1. Reads that originated from different genes correspond to distinct molecules, so if they share the same UMI then the UMI was sampled more than once (i.e. the UMI is not duplicated due to PCR).

2. While the number of sampled molecules $k$ is unknown, the number of distinct UMIs can be measured directly.

We say that a UMI that has been sampled more than once has a *collision* (Figure 3), and we denote the number of UMIs that appear in more than one gene by $x$ (see Figure 3). We denote the number of distinct UMIs sequenced, i.e. $|supp(U)|$, by $d$, and the number of distinct UMIs observed to be from a gene $g$ by $d_g$. We denote the number of sampled molecules originating from a gene $g$ by $k_g$. Note that $\sum_g d_g \geq d$.

We obtain method of moment estimates for the parameters $k, k_g$ and $n$ by relating them to $d_g, d$ and $x$. First, from equation (1) (see also Grün, Kester, and Oudenaarden 2014), we have that

$$d = n\left(1 - \left(1 - \frac{1}{n}\right)^k\right), \tag{2}$$

and at the gene level,

$$d_g = n\left(1 - \left(1 - \frac{1}{n}\right)^{k_g}\right). \tag{3}$$

Furthermore,

$$x = n\left(1 - \left(\left(1 - \frac{1}{n}\right)^k + \sum_g \left(1 - \left(1 - \frac{1}{n}\right)^{k_g}\right) \cdot \left(1 - \frac{1}{n}\right)^{k-k_g}\right)\right). \tag{4}$$

From equations (2) and (3) (see also Petukhov et al. 2018), we have that

$$\left(1 - \frac{1}{n}\right)^k = \frac{n-d}{n} \tag{5}$$

and at the gene level

$$\left(1 - \frac{1}{n}\right)^{k_g} = \frac{n-d_g}{n}. \tag{6}$$

Therefore, substituting equations (5), (6) into equation (4) we obtain

$$\begin{aligned} x &= n\left(1 - \left(\frac{n-d}{n} + \sum_g \left(1 - \frac{n-d_g}{n}\right) \cdot \left(\frac{n-d}{n-d_g}\right)\right)\right) & (7)\\ &= n\left(\frac{d}{n} - \frac{n-d}{n}\sum_g \left(\frac{d_g}{n-d_g}\right)\right) & (8)\\ &= d - (n-d)\sum_g \left(\frac{d_g}{n-d_g}\right). & (9) \end{aligned}$$

Since $d$, $d_g$ (for all $g$) and $x$ are known this equation can be used to estimate the effective number of UMIs $n$.

### 4 Estimating counts lost for each gene

Returning to equation (3), we see that

$$\hat{k_g} = \frac{ln\left(1 - \frac{d_g}{n}\right)}{ln\left(1 - \frac{1}{n}\right)}. \tag{10}$$

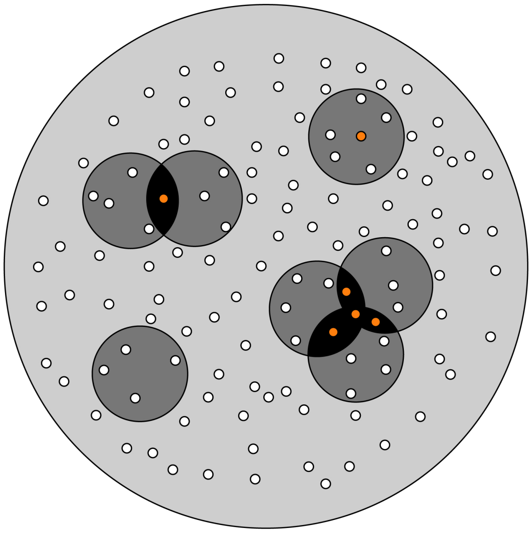

Figure 3: Collisions of UMIs. Each small white circle represents a distinct UMI. Each medium sized circle is a gene, and the enclosing circle is the set of all distinct UMIs. UMIs that have collided are shown in orange. Inter-gene collisions consist of UMIs present in two or more genes. An intra-gene collision is also shown.

With an estimate of $n$, and measurements of the quantities $d_g$, it is therefore possible to evaluate $k_g$, the number of molecules captured per gene. The loss of counts due to collapsing of UMIs by gene, is therefore given by

$$\hat{k_g} - d_g = \frac{ln\left(1 - \frac{d_g}{n}\right)}{ln\left(1 - \frac{1}{n}\right)} - d_g \quad (11)$$

$$\approx \frac{d_g(d_g - 1)}{2n + 1}. \quad (12)$$
